# Testing the predictive performance of comparative extinction risk models to support the global amphibian assessment

**DOI:** 10.1101/2023.02.08.526823

**Authors:** P.M. Lucas, M. Di Marco, V. Cazalis, J. Luedtke, K. Neam, M.H. Brown, P. Langhammer, G. Mancini, L. Santini

## Abstract

Assessing the extinction risk of species through the IUCN Red List is key to guiding conservation policies and reducing biodiversity loss. This process is resource-demanding, however, and requires a continuous update which becomes increasingly difficult as new species are added to the IUCN Red List. The use of automatic methods, such as comparative analyses to predict species extinction risk, can be an efficient alternative to maintaining up to date assessments. Using amphibians as a study group, we predict which species were more likely to change status, in order to suggest species that should be prioritized for reassessment. We used species traits, environmental variables, and proxies of climate and land-use change as predictors of the IUCN Red List category of species. We produced an ensemble prediction of IUCN Red List categories by combining four different model algorithms: Cumulative Link Models (CLM), phylogenetic Generalized Least Squares (PGLS), Random Forests (RF), Neural Networks (NN). By comparing IUCN Red List categories with the ensemble prediction, and accounting for uncertainty among model algorithms, we identified species that should be prioritized for future reassessments due to high prediction versus observation mismatch. We found that CLM and RF performed better than PGLS and NN, but there was not a clear best algorithm. The most important predicting variables across models were species range size, climate change, and landuse change. We propose ensemble modelling of extinction risk as a promising tool for prioritizing species for reassessment while accounting for inherent models’ uncertainty.

## 1. INTRODUCTION

Understanding how global change is affecting the extinction risk of species is key to guiding conservation planning and action (Pereira et al. 2010; Urban et al. 2016). The International Union for Conservation of Nature Red List of Threatened Species (RL) is responsible for monitoring the extinction risk of over 150,000 species globally (IUCN 2021), with experts assigning each species a category of extinction risk based on a set of quantitative criteria and thresholds (IUCN 2012). RL assessments require regular updates to ensure the information is sufficiently recent to inform and catalyze conservation action (IUCN 2012). However, insufficient resources are available to complete regular reassessments and expand its taxonomic coverage, and as a result, RL assessments face a risk of becoming outdated and not presenting a comprehensive picture of extinction risk (Cazalis et al. 2022; Rondinini et al. 2014). The long-term sustainability of the RL will depend on cost-efficient reassessment strategies (Rondinini et al. 2014), such as the identification of priority species for reassessment based on modelled probability of extinction risk.

Among terrestrial vertebrates, amphibians are the class with the highest proportion of threatened species, and they represent a particular challenge for extinction risk monitoring as the number of newly described species grows considerably every year (Ceballos et al. 2017; Stuart et al. 2004; Tapley et al. 2018) and many species have shown rapid declines (Sodhi et al. 2008; Wake and Vredenburg 2008). Climate change and habitat loss are deemed responsible of several drastic changes in amphibian conservation status in relatively short times. Examples are the Atacama Toad (*Rhinella atacamensis*) that was listed as Least Concern in 2010 and Vulnerable in 2015, and the Southern leopard frog (*Lithobates miadis*) that was classified as Vulnerable in 2004 and Critically Endangered in 2020.

Amphibians are characterized by traits that make them more prone to be negatively affected by global changes than other vertebrates, such as small geographic ranges, limited dispersal ability, dependence on water bodies, and limited thermoregulating abilities and sensitivity to evaporation (Duellman and Trueb 1994; Ficetola et al. 2015). However, while previous studies have shown that biological traits such as body size, geographic range size, or brood size play a key role in predicting amphibian RL extinction risk (Cardillo 2021; Cooper et al. 2008; Fontana et al. 2021; Pincheira-Donoso et al. 2021a; Sodhi et al. 2008), the effects of climate change remain poorly known. Developing robust tools that support the identification of species most in need of reassessment is highly valuable for informing global extinction risk monitoring strategies (Cazalis et al. 2022).

Comparative extinction risk models that relate RL categories with a number of extrinsic or intrinsic drivers are one of the tools that can support RL assessments with limited resources (Cazalis et al. 2022). RLComparative analyses of extinction risk have typically relied on single model algorithms (Bland et al. 2015; Di Marco et al. 2014; Wieringa 2022; Zizka et al. 2022; Zizka et al. 2021). When several models are used, a best model is generally selected based on individual predictive performance (Bland et al. 2015). However, a single best model may not result in the best predictions, relying on several good performing models may be a better strategy whilst allowing to inform on predictive uncertainty (Araujo and New 2007).

Here we use a combination of comparative extinction risk models to identify amphibian species that should be prioritized for reassessment, based on their potential RL category. Specifically, we evaluate the effects of species traits, environmental variables, and global changes (including climate and land use change) on species extinction risk. We consider several different model algorithms and assess their relative predictive power and produce ensemble predictions that account for uncertainty expressed as inconsistency among model predictions to inform the prioritization of RL reassessments.

## 2. MATERIAL AND METHODS

### 2.1. Predictors of extinction risk

#### 2.1.1. Species distributions and spatial variables

We retrieved geographic range polygons for all amphibian species from the RL dataset (IUCN 2021). We considered range polygons classified as native or reintroduced in origin, with extant or probably extant presence (IUCN 2018). We projected the selected polygons using the Lambert Cylindrical Equal Area (CEA) projection and then calculated a series of spatial variables within each amphibian distribution: 38 variables describing the mean and standard deviation of the 19 bioclimatic variables (Karger et al. 2017; Table A1); 19 variables describing the change in bioclimatic variables between 1965–1994 and 2005–2014 (Karger et al. 2017, 2018; Table A1 and Supplementary methods); 4 variables describing land cover/land cover change: percentage of urban areas, percentage of agriculture areas, change in percentage of the urban areas, and change in percentage of agriculture areas (ESA 2021; Table A2); 3 variables describing anthropogenic pressures: mean value for accessibility (Weiss et al 2018; Table A2), mean human population density, and change in human population density (NASA 2018; Table A2); 10 variables describing the biogeographical realms, the most representative realm, the sum of number of realms where the species occurs, and a variable for each realm indicating whether the species is present/absent (Olson et al. 2001; Table A1); and 4 descriptors of the geographic range of the species, including range area and spatial configuration (Lucas et al 2019; Tables A1 and A2, Supplementary methods).

#### 2.1.2. Biological traits

For each species, we compiled information from existing datasets (IUCN 2021; Oliveira et al. 2017; Pincheira-Donoso et al. 2021a; Pincheira-Donoso et al. 2021b) for 32 biological traits such as nesting site, microhabitat, parity mode, diet, daily/seasonal activity pattern, number of habitats, and body size (Table A1). To account for species phylogeny, we extracted the first six eigenvectors from the phylogeny in Jetz et al. 2018, which explained 96% of the total variance (Table A3). Due to the continuous and extensive changes in amphibian taxonomy at species level, different datasets use varying taxonomic nomenclature. To correct for this discrepancy, we used “AmphiNom” package (Liedtke 2019) to combine the different datasets, by first obtaining all listed synonyms per species from Amphibian Species of the World (i.e., the taxonomic authority for amphibians on the RL).

To fill data gaps in the compiled biological trait dataset, we applied a data imputation procedure. We used the biological trait dataset along with the taxonomic order, six eigenvectors describing the phylogeny, the variables describing the biogeographical realms, range area, and the climate variables (see spatial variables section above) -- a total of 88 variables for each species -- to impute missing life-history data (Table A1). We scaled all numerical variables to mean = 0 and standard deviation = 1, and then used the ‘missForest’ package in R to infer the missing values (Stekhoven and Bühlmann 2011). We calculated the Normalized Root Mean Square Error (NRMSE) for imputed data, and discarded trait variables with an NRMSE > 40% and a proportion of missing data > 40% (Penone et al. 2014; Table A4 and Figure A1).

#### 2.1.3. Data preparation

We used spatial information, biological traits, and phylogeny as predictor variables to model the extinction risk of amphibians. To reduce the overfitting of the models and to have a dataset manageable to fit and validate all model algorithms, we pre-selected and/or modified some of the available variables (Supplementary methods, Tables A2-A8 and Figure A1). For climate variables, and separately for climate change variables, we applied a principal component analysis (PCA) and selected the first four components that explained 74% of the variance for climate and the first five components that explained 93% for climate change (Supplementary methods, Tables A5 and A6). For species phylogeny, we used the first four eigenvectors from the phylogeny which explained at least 93% of the variance (Table A4). For realm variables, we selected the *Realm* variable which describes the dominant biogeographical realm for each species (Supplementary methods and Table A2). For variables describing species habitat specialization, we classified species as generalist (living in more than one habitat) or not, and as occurring in forest or not (Supplementary methods and Tables A2 and A7), and we grouped the variable *Microhabitat* (Table A8).

We divided the dataset by taxonomic order – Anura, Caudata, and Gymnophiona – and scaled all predictor variables and eliminated the colinear variables with variance inflation factor (VIF) > 10 (Table A9).

### 2.2. Modelling extinction risk

We generated separate models for each amphibian order because we expected correlates of extinction risk to differ among them (González-del-Pliego et al. 2019). Following Cazalis et al. (2022), who reviewed recent efforts in comparative extinction risk models, we applied four commonly used algorithms: cumulative link models (CLM), phylogenetic generalized least squares models (PGLS), random forest models (RF) and artificial neural networks models (NN). For predictor variables, we used numeric and Boolean predictors in CLM and NN, and numeric and factor predictors for RF and PGLS. As a response variable, we used five RL categories: Least Concern, Near Threatened, Vulnerable, Endangered, Critically Endangered. In CLM models we used RL categories as ordinal factor variables. In RF models we used RL categories as a factor variable. For NN, and PGLS algorithms RL categories were transformed to a numerical variable: Least Concern = 1, Near Threatened = 2, Vulnerable = 3, Endangered = 4, Critically Endangered = 5. Notice that RF and NN algorithms, can use either numerical or factor response variables, we chose the response variable based on the best performance of preliminary analysis (results not shown). For each model algorithm we used different procedures for model/variable selection and the selection of parameters/hyperparameters (Supplementary methods and Table A9). To make predictions comparable and to allow for the use of the same validation measures, all model predictions were transformed to an integer variable. For CLM we selected the integer value with the highest probability in the prediction. For PGLS and NN the continuous predictions (*predictions*) were transformed to numerical values, as follows: *predictions* < 1.5 = 1, 1.5 ≤ *predictions* < 2.5 = 2, 2.5 ≤ *predictions* < 3.5 = 3, 3.5 ≤ *predictions* < 4.5 = 4, 4.5 ≤ *predictions* = 5. For RF, factorial predictions of RL categories were transformed to a numerical variable: Least Concern = 1, Near Threatened = 2, Vulnerable = 3, Endangered = 4, Critically Endangered = 5.

Comparative extinction risk analyses that aim to identify the drivers of extinction risk (Lucas et al. 2019), usually exclude species classified under criterion B to avoid a circularity. Indeed, the extent of occurrence and the area of occupancy that are used for application of criteria B1 and B2, respectively, are highly correlated with geographic range size that is typically included as model predictor. Such circularity can lead to an overestimation of the importance of range size, obscuring the role of other important drivers. Instead, when comparative extinction risk analyses have a predictive goal, species classified under criterion B are not excluded in order to generate the best possible predictions (see e.g. Zizka et al. 2021, Caetano et al. 2022). As the main goal of our models is to obtain the best predictions of extinction risk to assess possible mismatches with official RL assessments, rather than evaluating the actual importance of range size in comparison to other variables, we retained species assessed under criterion B.

### 2.3. Variable importance

We calculated the variable importance (*VI*) using different methodologies. For CLM and PGLS, we used the scaled coefficients for each variable:

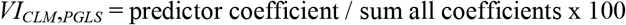

For RF models, we used the mean decrease accuracy (MDA) which is based on how much the accuracy decreases when the variable is excluded:

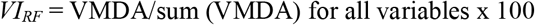

For NN, we used the implemented method by Gedeon (Gedeon 1997) which considers the weights connecting the input features to the first two hidden layers:

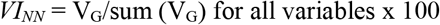

Using all model algorithms, we calculated the average importance of each variable for each amphibian order, and combining all models we calculated the average importance of each variable for the class Amphibia.

### 2.4. Model validation

We evaluated each model’s performance using taxonomic block validation, iteratively extracting one family from the dataset for testing and then fitting the model on the remaining families (Roberts et al. 2017). This approach is more robust to assess models’ predictive performance compared to the random cross-validation used in several previous studies (González-del-Pliego et al. 2019), as random cross validation tends to produce over-optimistic results due to the autocorrelation between phylogenetically close species. However, to compare our results to previous comparative extinction risk models on amphibians, we also ran a classical random cross-validation extracting 10% of the species in each run and repeating the operation 10 times. For each predicted set we validated the models at two different levels. First, the models were validated at the RL category level by calculating the overall accuracy (rate of correct classification) and the mean classification error (absolute value of the difference between predicted and observed categories) for all categories and the overall accuracy, the sensitivity (rate of correct classification of threatened species), the specificity (rate of correct classification of non-threatened species), and the true skill statistic (TSS = Specificity + Sensitivity −1). Additionally, for each category, and to ensure comparability with previous studies of extinction risk, we also aggregated the predictions into binary classes of non-threatened (LC and NT) versus threatened (VU, EN, CR), and calculated the overall accuracy, sensitivity, specificity, and TSS.

### 2.5. Ensemble prediction and species prioritization

We used an ensemble forecasting approach for predicting extinction risk (Araujo and New 2007). To calculate the ensemble prediction, we combined the predictions of all models that achieved an acceptable predictive performance. We considered predictive performance as “acceptable” when mean error was below 1.00 during block validation (i.e., less than one category mismatch). Using these subsets of models, we calculated the ensemble prediction for each species as the mean of the predictions from the individual models. In addition, we calculated the standard deviation of extinction risk prediction among the subset of models.

Then we calculated the *Difference in Extinction risk* (*DE*) using the ensemble prediction and the current RL category of the species.

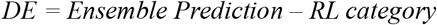

A positive difference represents species that are predicted to be more threatened than the published RL category (i.e. *overpredicted species*). Conversely, a negative difference indicates species that are predicted to be less threatened than the published RL category (i.e. *underpredicted species*).

To account for the variability across model predictions, we also calculated the *Scaled Standard Deviation* (*SSD*) as the coefficient of each SD by the maximum SD for each taxonomic order, which resulted in a scaled value between 0 and 1. Then we used the *DE* and the *SSD* to calculate a *Species Prioritization Index (SPI*). To differentiate *overpredicted species* and *underpredicted species*, we applied the index separately to these two groups obtaining two *SPIs:* (1) *Species Prioritization Index Overpredicted (SPI_O_*) for *overpredicted species*, species for which their RL category shows a lower extinction risk category than our predicted category; and (2) *Species Prioritization Index Underpredicted (SPI_O_*) for *underpredicted species*, species for which their RL category is higher than our predicted category.

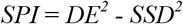

High *SPI* values are represented by species with high *DE* and with low values of *SSD* among the predictions of the algorithms included. High *SPI_O_* values indicate species that should be particularly prioritized for reassessment because they are predicted to be more threatened than currently assessed (according to the ensemble prediction). High *SPI_U_* values instead indicate species that are predicted to be less threatened than currently assessed.

To visualize the spatial pattern of assessment priorities, we intersected species range maps with a grid of 100 x 100 km near the equator (Harfoot et al. 2021) using ArcMap 10.3 (ESRI 2008), and calculated for each grid cell the average *SPI_O_;* and the average *SPI_U_*. In addition, to determine if there was a taxonomic bias in the *SPI* values, we calculated for each family the *SPI_O_*; and *SPI_U_*.

## 3. RESULTS

### 3.1. Predictors of extinction risk in amphibians

The most important variables explaining extinction risk for all amphibians were geographic range area (*Range_Area*), followed by the realm in which the species is present (*Realm*), climate variables (*Climate_2, Climate_4* and *Climate_1*), habitat specialisation variables (*Habitat_generalist_forest*), levels of human presence (e.g., *Urbanization* and *Accessibility*), climate change (*Climate_Change_5*), and the microhabitat where the species is found (*Microhabitat;* Figure 1 and Table A10). *Range_area* and *Climate_Change_5* were consistently important across the three amphibian orders, but some variables were specifically important for certain taxa, such as *Accessibility* for Anura or *Climate_2* for Gymnophiona. As expected, *Range_area* was negatively associated with extinction risk (i.e., species with larger ranges are less likely to be threatened), and lower *Accessibility* (less i.e., less travel time to cities, less distance from cities) and higher *Urbanization* were associated with higher extinction risk (Tables A10-A16 and Figures A2-A7).

**Fig. 1.**
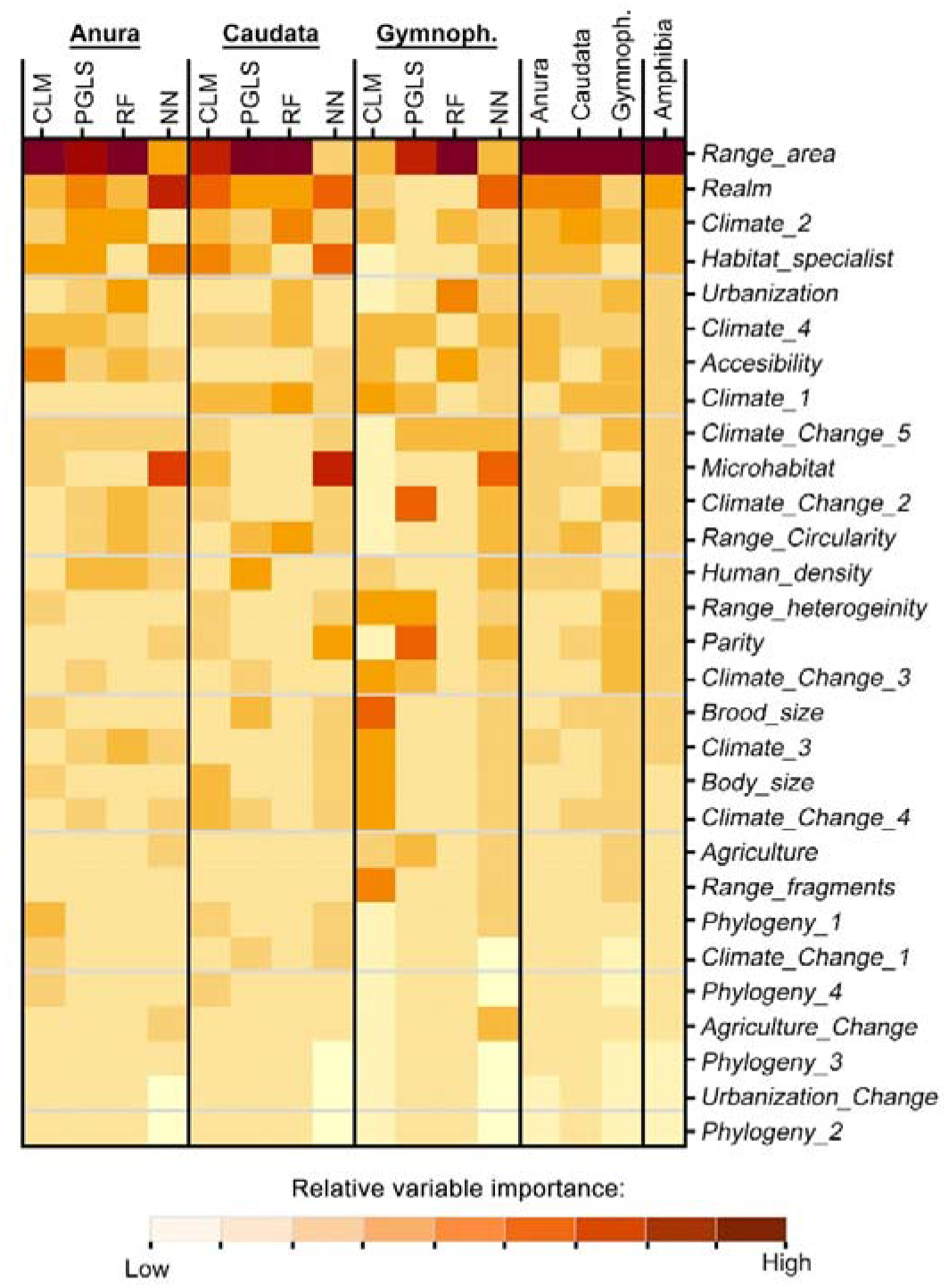
Variable importance to predict extinction risk of amphibians according to the four models (Cumulative Link Models [CLM], Random Forest [RF], Phylogenetic Generalized Least Square models [PGLS], Neural Network [NN]) per taxonomic order (Anura, Caudata, and Gymnophiona). The four rightmost columns Anura, Caudata, Gymnophiona, and Amphibia indicate the average importance per taxonomic group. Variable importance has been scaled within each column so values of variable importance are relative within each column and not comparable between different columns. Original (non-scaled) values are available in Table A9.

### 3.2. Model validation

Block family validation showed an accuracy averaged among categories (independently of the number of species in each category) and all models of 0.82 ± 0.09 (Mean ± SD). Gymnophionans had the highest accuracy (0.85 ± 0.12), followed by anurans (0.81 ± 0.08) and caudatans (0.79 ± 0.07). RF and CLM had the highest accuracy values, with 0.87 ± 0.06 and 0.83 ± 0.08 respectively (Fig. 2a, Tables A17 and A18).

**Fig. 2.**
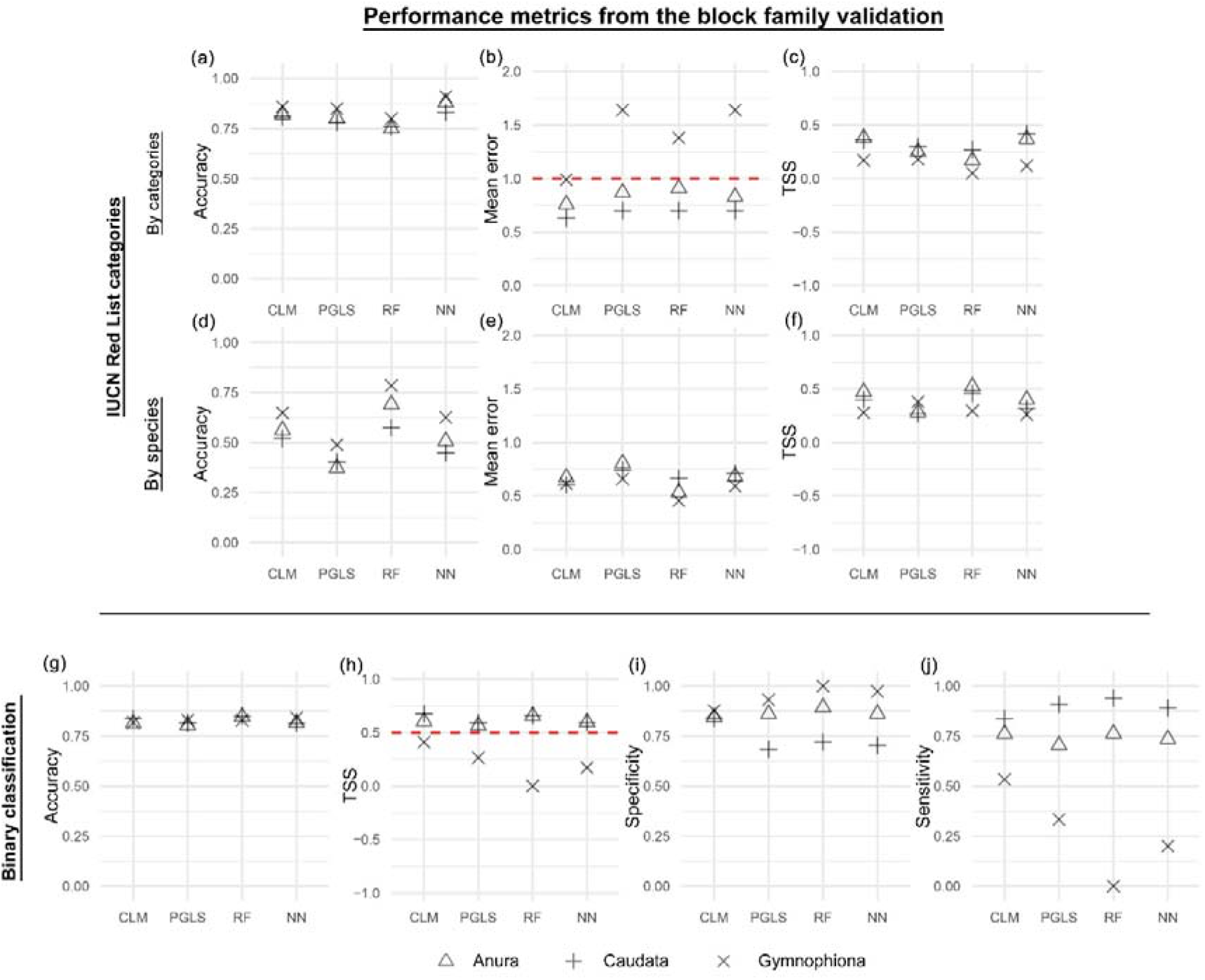
Performance metrics from the family block validation for four model algorithms (Cumulative Link Models [CLM], Random Forest [RF], Phylogenetic Generalized Least Square models [PGLS], Neural Network [NN]) considering RL categories for each of the three amphibian orders (a-f). We report the accuracy (rate of correct classification in all categories), mean error (absolute value of the difference between predicted and current RL categories) and true skill statistic (TSS = Specificity + Sensitivity −1) averaged among RL categories (independently of the number of species in each category; top row) and by the number of species (middle row). Mean error averaged among categories (b) was used to exclude models for the ensemble prediction when the value was ≥1.00 (red horizontal line indicates mean error = 1.00). Values averaged among species (d-f) are reported to compare with classical performance metrics reported in comparative studies of extinction risk. Performance metrics from the family block validation considering binary classification (threatened/non-threatened) risk categories (g-j). Accuracy (rate of correct classification), true skill statistic (TSS = Specificity + Sensitivity − 1), specificity (rate of correct classification of non-threatened species), and sensitivity (rate of correct classification of threatened species) are reported for each amphibian order and each model algorithm. For the TSS plot (h), the horizontal dashed line (TSS = 0.5) indicates the value at which models are considered good.

The TSS averaged across categories showed a mean value of 0.25 ± 0.22 for all models. By order, Caudata models showed the best TSS values (0.34 ± 0.18), followed by Anura (0.29 ± 0.20) and Gymnophiona, which showed the lowest TSS values (0.13 ± 0.22) among all models. Among the models, CLM and RF models showed the best TSS results with 0.30 ± 0.2212 and 0.30 ± 0.25 respectively, followed by NN (0.25 ± 0.15) and PGLS (0.16 ± 0.22; Fig. 2c).

The mean error showed on average a value of 0.98 ± 0.73 among all categories and models, indicating that, on average, the mean error was of less than one category. However, it showed important differences among the three orders; Caudata showed the lowest mean error (0.68 ± 0.26), followed by Anura (0.84 ± 0.32) and Gymnophiona (1.41 ± 1.07). CLM showed the lowest error (0.79 ± 0.32) among models (Fig. 2b).

When we averaged the values by the number of species (instead of by categories), our independent family block validation showed an average accuracy of 0.55 ± 0.12 among all models (Fig. 2d). RF showed the best results (0.68 ± 0.11), followed by CLM (0.58 ± 0.07), NN (0.53 ± 0.09) and PGLS (0.42 ± 0.06). Gymnophiona showed the best accuracy (0.64 ± 0.12) followed by Anura (0.53 ± 0.13) and Caudata (0.49 ± 0.08, Table A19).

The TSS averaged across species showed a mean value of 0.36 ± 0.09 for all models. By order, Anura models showed the best TSS values (0.42 ± 0.11), followed by Caudata (0.36 ± 0.08) and Gymnophiona, which showed the lowest TSS values (0.30 ± 0.05) among all models. Among the models, RF models showed the best TSS results with a mean of 0.42 ± 0.12, followed by CLM (0.38 ± 0.10), PGLS (0.31 ± 0.06) and NN (0.33 ± 0.07; Fig. 2f).

The mean error showed an average value among all models of 0.64 ± 0.094. RF algorithms showed on average the lowest mean error (0.55 ± 0.11), followed by CLM (0.63 ± 0.04), NN (0.66 ± 0.06) and PGLS (0.74 ± 0.07). The mean error was lowest for Gymnophiona (0.58 ± 0.09), followed by Anura (0.67 ± 0.11) and Caudata (0.68 ± 0.06) (Fig. 2e).

When considering family block validation for binary classification (threatened/non-threatened species), we obtained an average accuracy of 0.83 ± 0.02 across all models (Fig. 2g-j; Table A19). Accuracy values were very similar among the four model algorithms (SD = 0.01), with 0.84 ± 0.01 in RF, 0.82 ± 0.01 in CLM and PGLS, and 0.82 ± 0.01 in NN. Average TSS across all models was 0.48 ± 0.22. There was a low variability in TSS among different model algorithms (SD = 0.06) and high variability among the orders (SD = 0.24). Caudatans and anurans had on average a similar TSS (0.63 ± 0.04 and 0.60 ± 0.04 respectively), while gymnophionans models showed lower values (mean TSS = 0.21 ± 0.17). Among all algorithms tested, CLM resulted in the highest TSS (TSS = 0.56 ± 0.14). In general, all models showed a good balance for specificity and sensitivity values, except for PGLS, RF and NN models for gymnophionans. CLM models showed on average showed on average the least absolute difference between specificity and sensitivity in the three orders (Fig. 2g-j).

Random cross-validation led to substantially better estimates. Mean error by category averaged among categories = 0.72 ± 0.39; average TSS for binary classification among all models of 0.56 ± 0.23, TSS = 0.65 ± 0.06 for anurans, TSS = 0.72 ± 0.06 for caudatans, and TSS = 0.31 ± 0.24 for gymnophionans. (Tables A21-A24 and Figures A8 and A9).

### 3.3. Ensemble prediction and species prioritization

Based on the mean error averaged among categories and using the RL categories as a response variable (see above), we excluded three models with unacceptable errors (≥1.00; Fig. 2a, Figure A7) to calculate the ensemble prediction: PGLS, RF and NN models for gymnophionans. The ensemble prediction indicates that 40.46% of species (n=2,300) might be threatened with extinction, compared to 38.18% species (n=2,170) assessed as such (Table A25). According to our predictions 40% (n=2,274) species are underpredicted (*DE*>0) and 30% (n=1,697) species are overpredicted (*DE*<0), with 30% (n=1,713) species having the same predicted values as the current RL category (Fig. 3 and Figure A10). However, most of *DE* values are relatively small, and only 3.47% (n=197) have a *DE* higher than 2 or smaller than −2.

**Fig. 3.**
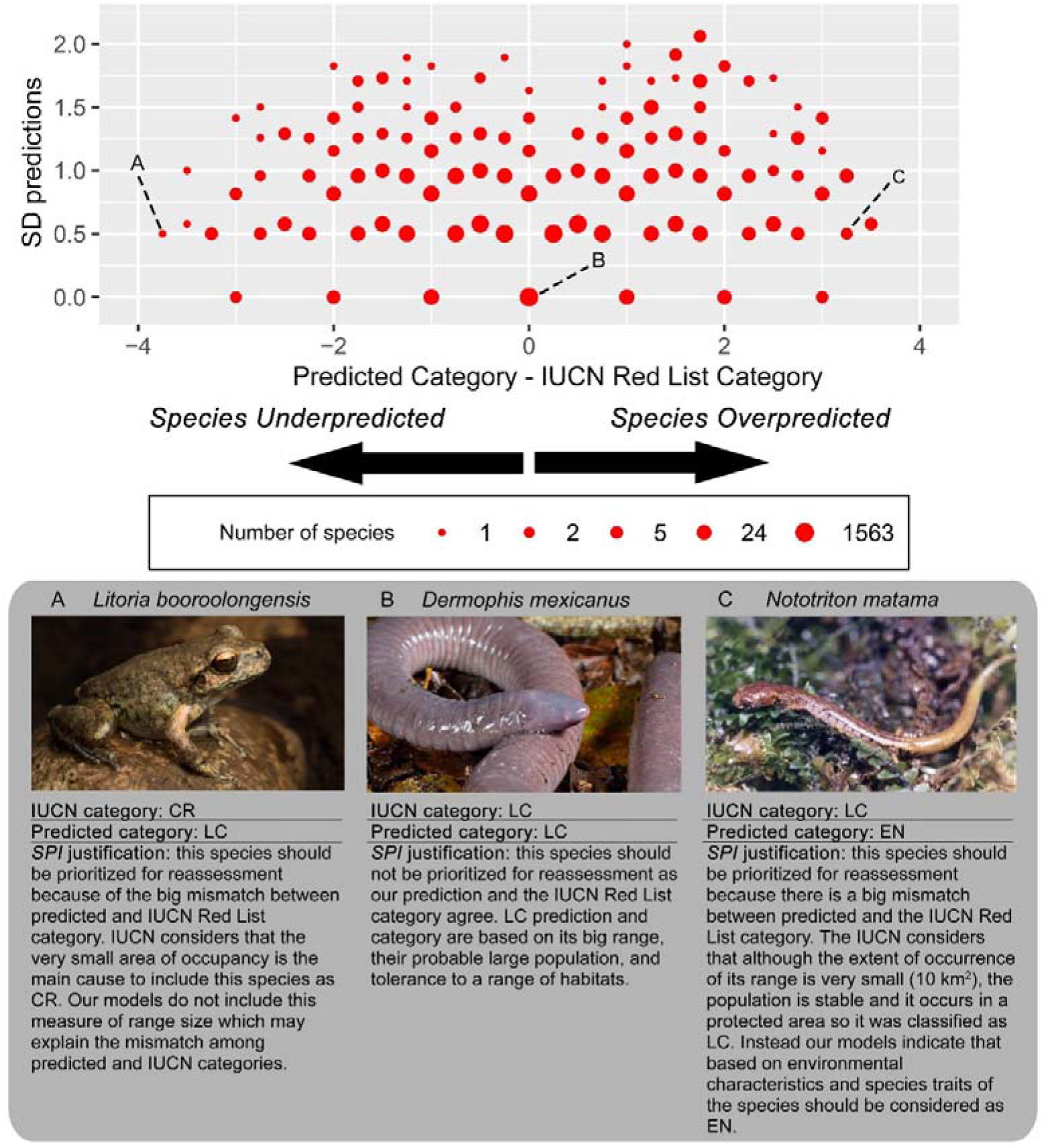
Distribution of the number of species by values of *DE* (Predicted Category - RL Category) and standard deviation (SD) of the predictions. Species with only one model with TSS > 0.4 (Gymnophiona) are plotted as with SD = 0. Photos A by Matthew Clancy, B by José G. Martínez-Fonseca and C by Eduardo Boza-Oviedo.

The distribution of *SPI* was positively skewed, with values ranging from 14.00 indicating a high priority for reassessment to −0.78 which indicates a low reassessment priority (Table A25 and Figure A10). *Overpredicted species* rate (corrected by the number of species on the same grid to control for the species richness) is concentrated mainly in Central America, Andes region, eastern and western coast of the USA, and southeast Asia. *Underpredicted species* rate (corrected by the number of species on the same grid to control for the species richness) is concentrated mainly in Central America, Andes region, west coast of the USA, southwest Europe, southeast Asia and southeast Australia (Figure A11). *Leiopelmatidae* and *Rhinodermatidae* families had substantially higher *SPI_U_* values than the average *SPI*, while *Rhinatrematidae* and *Ichthyophidae* families had substantially higher *SPI_O_* values than the average *SPI* (Figure A12).

## 4. DISCUSSION

Here we presented an ensemble of comparative predictive models to assess their potential to prioritize amphibian species for reassessment on the RL. We compared several predictive models which exhibit diverse predictive power, and while all models generally showed good predictive performance, no single model algorithm worked best. Therefore, we propose an approach to predicting species extinction risk that combines the predictions of models with sufficient predictive power while accounting for their consistency. This approach prioritizes species whose predicted RL category differs the most from its current category on the RL, and for which predictions are consistent among models.

### 4.1. Predictors of extinction risk in amphibians

The most important variable across our models was range size. This variable is commonly used in comparative extinction risk analyses (Chichorro et al. 2019) and often results in a negative relationship with extinction risk (including amphibians: Gonzalez del Pliego et al. 2019, Sodhi et al. 2008, Cooper et al. 2008), which is not surprising given the large proportion of species classified as threatened under Criterion B (IUCN 2012). Other range-related variables were also important, particularly the realm where species occur, echoing studies that found extinction risk is highly structured in amphibians (González-del-Pliego et al. 2019), and to some extent the shape of the range as previously reported for terrestrial vertebrates (Lucas et al. 2019). Among human pressure variables, urbanization and accessibility were the most important factors, probably providing proxies for population declines due to human activities (Cardillo 2021; González-del-Pliego et al. 2019; Sodhi et al. 2008). Climate and climate change covariates were also strong predictors of extinction risk. This is a novel and important result because climate change effects are emerging as a serious threat to this group of species (Cohen et al. 2019; Lertzman-Lepofsky et al. 2020; Miller et al. 2018), but the lack of information on species-specific effects of climate change make it difficult for RL assessors to take climate change into account (Cazalis et al. 2022; Foden and Young). Our results could help assessors in identifying those species that may be suffering the most from climate change.

Among traits, habitat specialization was the fourth most important covariate predicting extinction risk. We observed lower risk of extinction for generalist species living in forest compared to generalists not living in forest, which may be associated with the potential of forests to buffer the negative effects of climate change (De Frenne et al. 2021; De Frenne et al. 2019; Ramalho et al. 2021). Conversely, body size and reproductive traits were poor predictors of risk in this study. Phylogeny has previously been shown to be an important predictor of amphibian extinction risk (González-del-Pliego et al. 2019), mainly because the different traits and factors leading to increased extinction risk are highly structured phylogenetically. In contrast, phylogeny was a very weak predictor in our analysis, and PGLS did not perform better than other models, suggesting that phylogeny may be a proxy of other trait covariates used in our analysis.

### 4.2. Performance of the different model algorithms

No modelling approach consistently outperformed the others, but several approaches produced good results for the two most speciose taxonomic orders (Anura and Caudata). The overall accuracy of our models at the category level was high with predictions diverging by <1 Red List category from the published assessments on average. It should be considered, however, that when predictions are made for species in families not included in the training dataset, our family-block validation likely tends to under-estimate the models’ true predictive ability. As expected, random cross-validation led to substantially better estimates, which are comparable to previous comparative extinction risk analysis of mammals (Bland et al. 2015), reptiles (Caetano et al. 2022), and plants (Zizka et al. 2021), and are higher than the latest analysis on amphibians (González-del-Pliego et al. 2019).

The performance of the different algorithms showed important variations among the three amphibian orders. Random Forests and Neural Network models performed better for anurans and caudatans, suggesting that sample size may be a limiting factor in these complex models (Tange et al. 2017; Vabalas et al. 2019). This may explain why simpler models, such as the CLM, outperformed the other model algorithms for gymnophionans.

Our results show that extinction risk can be predicted with good performance for the vast majority of amphibians, a group that was comprehensively assessed for the RL in 2004 (Stuart et al. 2004) but has since proven difficult to maintain up-to-date assessments due to the high rate of new species descriptions, rapid response to environmental stressors, and limited resources available for reassessment. The use of multiple model algorithms allowed us to select the best performing models, overall improving performance compared to the use of a single algorithm. In addition, producing several equally valid predictions of extinction risk provide complementary information of model uncertainty that can be used to weight the predictions to identify reassessment priorities.

### 4.3. Applications, limitations and future steps

Spatial and taxonomic patterns in *SPI* can be informative for reassessment prioritization, as well as interpretation of models’ limitations. A mismatch between observed and predicted categories can arise from multiple mechanisms.

An overprediction of extinction risk can be informative for identifying species that may have a higher extinction risk than the current assessment, even though not currently threatened. This may be the case of many Least Concern or Near Threatened species which may be prioritized for reassessment. This can result from the use of ancillary information in the model (e.g., climate change, land-use change, etc.) that was not directly used in the assessment process, either because it was unavailable to the assessors, or they were unaware of the information. In addition, overpredicted species that are currently Least Concern or Near Threatened could be monitored for emerging threats that would qualify them for a threatened category. Over-prediction can also be a modelling artefact, which disregards the complex and composite nature of Red List assessments that typically require multiple subcriteria to be met in order for a species to be considered threatened (Di Marco 2022), e.g. restricted range size and continuing population decline.

In contrast, underpredictions are likely a derivative, in most cases, of model simplification. Among the most pressing threats to amphibians are pathogens and invasive species (Scheele et al. 2019; Stuart et al. 2004; Stuart et al. 2008), which could not be included explicitly in our models and might explain those mismatches and likely explain the majority of underpredictions. Even less likely, it is also possible that underpredictions may point to errors in RL assessments that may need revision.

Finally, mismatches may also indicate possible inconsistencies in the assessments of different families or genera, or different regions of the world assessed by different specialist groups. Such inconsistencies can, for example, arise from the predominant application of certain criteria or data types used (or not used) in the assessments. All in all, the SPI values for species should be assessed on a case-by-case basis by experts to help define future reassessment priorities. Assessors may undoubtedly have additional factors to consider in the prioritization, which may span from available funding for certain regions or taxonomic groups, to groups that have recently undergone taxonomic revision. In these cases, SPI can be used to prioritize pre-selected groups of species based on other criteria, in order to optimise the efforts required to maintain up to date RL assessments under limited available resources (Cazalis et al. 2022; Rondinini et al. 2014).

Extinction risk models are still unable to replicate the red listing process, but can be a powerful tool to assist and optimize the work of assessors by pinpointing species potentially in need of reassessment. Future steps to further improve these models and their usability include the reevaluation of predictions based on new assessments (Di Marco 2022), and the identification of mechanisms that led to a mismatch between predicted and observed RL categories to identify inherent biases in either the modelling process (e.g. omission of a relevant variable or inclusion of a misleading one) or the assessment process (e.g. omission of a relevant information, such as climate change). Incorporating disease and future climate change scenarios, as well as real-time threat data (e.g., deforestation alerts) will also be critical next steps in refining the prioritization process for amphibian reassessments.

## AUTHOR CONTRIBUTIONS

L.S. and M.D.M. conceived the study; P.M.L. led the analysis of the data under the supervision of L.S., M.D.M., V.C., and G.M; P.M.L. interpreted the results with contributions from L.S., M.D.M., V.C., J.L., K.N. and G.M; P.M.L. led the writing of the manuscript with inputs from L.S., M.D.M., V.C., J.L., K.N., M.H.B., P.F.L. and G.M.

## ACKNOWLEDGMENTS

We are sincerely grateful to the working group sRedList for their helpful comments on this manuscript. We also thank the funding to sRedList by sDiv, the Synthesis Centre of the German Centre for Integrative Biodiversity Research (iDiv) Halle-Jena-Leipzig, funded by the German Research Foundation (FZT 118, 202548816), which also funded the position for V.Ca.. L.S. acknowledges support from the MUR Rita Levi Montalcini program. We thank Re:wild and Microsoft Corporation (Grant number: 5313.006-AI4E) who provided funding for the position of P.M.L, and Sapienza University of Rome who funded this work (AR22117A859F7D58). We acknowledge to Matthew Clancy, José G. Martínez-Fonseca and Eduardo Boza-Oviedo for their photos used in Figure 3.

## Notes

### Competing Interest Statement

The authors have declared no competing interest.

## REFERENCES

Amado, T.F., Martinez, P.A., Pincheira-Donoso, D., Olalla-Tárraga, M.Á., 2021. Body size distributions of anurans are explained by diversification rates and the environment. Global Ecology and Biogeography 30, 154–164.

Araujo, M.B., New, M., 2007. Ensemble forecasting of species distributions. Trends in Ecology & Evolution 22, 42–47.

Bland, L.M., Collen, B., Orme, C.D.L., Bielby, J., 2015. Predicting the conservation status of data-deficient species. Conservation Biology 29, 250–259.

Caetano, G.H.d.O., Chapple, D.G., Grenyer, R., Raz, T., Rosenblatt, J., Tingley, R., Böhm, M., Meiri, S., Roll, U., 2022. Automated assessment reveals that the extinction risk of reptiles is widely underestimated across space and phylogeny. PLoS Biology 20, e3001544.

Cardillo, M., 2021. Clarifying the relationship between body size and extinction risk in amphibians by complete mapping of model space. Proceedings of the Royal Society B: Biological Sciences 288, 20203011.

Cazalis, V., Di Marco, M., Butchart, S.H.M., Akçakaya, H.R., González-Suárez, M., Meyer, C., Clausnitzer, V., Böhm, M., Zizka, A., Cardoso, P., Schipper, A.M., Bachman, S.P., Young, B.E., Hoffmann, M., Benítez-López, A., Lucas, P.M., Pettorelli, N., Patoine, G., Pacifici, M., Jörger-Hickfang, T., Brooks, T.M., Rondinini, C., Hill, S.L.L., Visconti, P., Santini, L., 2022. Bridging the research-implementation gap in IUCN Red List assessments. Trends in Ecology & Evolution 37, 359–370.

Ceballos, G., Ehrlich, P.R., of the, D.-R., 2017. Biological annihilation via the ongoing sixth mass extinction signaled by vertebrate population losses and declines. Proceedings of the….

Chichorro, F., Juslén, A., Cardoso, P., 2019. A review of the relation between species traits and extinction risk. Biological Conservation 237, 220–229.

Cohen, J.M., Civitello, D.J., Venesky, M.D., McMahon, T.A., Rohr, J.R., 2019. An interaction between climate change and infectious disease drove widespread amphibian declines. Global Change Biology 25, 927–937.

Cooper, N., Bielby, J., Thomas, G.H., Purvis, A., 2008. Macroecology and extinction risk correlates of frogs. Global Ecology and Biogeography 17, 211–221.

De Frenne, P., Lenoir, J., Luoto, M., Scheffers, B.R., Zellweger, F., Aalto, J., Ashcroft, M.B., Christiansen, D.M., Decocq, G., De Pauw, K., Govaert, S., Greiser, C., Gril, E., Hampe, A., Jucker, T., Klinges, D.H., Koelemeijer, I.A., Lembrechts, J.J., Marrec, R., Meeussen, C., Ogée, J., Tyystjärvi, V., Vangansbeke, P., Hylander, K., 2021. Forest microclimates and climate change: Importance, drivers and future research agenda. Global Change Biology 27, 2279–2297.

De Frenne, P., Zellweger, F., Rodríguez-Sánchez, F., Scheffers, B.R., Hylander, K., Luoto, M., Vellend, M., Verheyen, K., Lenoir, J., 2019. Global buffering of temperatures under forest canopies. Nature Ecology & Evolution 3, 744–749.

Di Marco, M., 2022. Reptile research shows new avenues and old challenges for extinction risk modelling. PLoS Biology 20, e3001719.

Di Marco, M., Buchanan, G.M., Szantoi, Z., Holmgren, M., Grottolo Marasini, G., Gross, D., Tranquilli, S., Boitani, L., Rondinini, C., 2014. Drivers of extinction risk in African mammals: the interplay of distribution state, human pressure, conservation response and species biology. Philosophical Transactions of the Royal Society B: Biological Sciences 369, 20130198.

Duellman, W.E., Trueb, L., 1994. Biology of amphibians. Johns Hopkins University Press, Baltimore.

ESA, 2021. Land cover classification gridded maps from 1992 to present derived from satellite observations version 2.0 and 2.1.

ESRI, 2008. ArcMap 9.3 Redlands, CA: Environmental Systems Research Institute.

Ficetola, G.F., Rondinini, C., Bonardi, A., Baisero, D., Padoa-Schioppa, E., 2015. Habitat availability for amphibians and extinction threat: a global analysis. Diversity and Distributions 21, 302–311.

Foden, W.B., Young, B.E., IUCN SSC Guidelines for Assessing Species’ Vulnerability to Climate Change. Version 1.0. Occasional Paper of the IUCN Species Survival Commission No. 59 IUCN Species Survival Commission, Cambridge, UK and Gland, Switzerland.

Fontana, R.B., Furtado, R., Zanella, N., Debastiani, V.J., Hartz, S.M., 2021. Linking ecological traits to extinction risk: Analysis of a Neotropical anuran database. Biological Conservation 264, 109390.

Gedeon, T.D., 1997. Data mining of inputs: analysing magnitude and functional measures. International Journal of Neural Systems 8, 209–218.

González-del-Pliego, P., Freckleton, R.P., Edwards, D.P., Koo, M.S., Scheffers, B.R., Pyron, R.A., Jetz, W., 2019. Phylogenetic and Trait-Based Prediction of Extinction Risk for Data-Deficient Amphibians. Current Biology 29, 1557–1563.e1553.

Harfoot, M.B.J., Johnston, A., Balmford, A., Burgess, N.D., Butchart, S.H.M., Dias, M.P., Hazin, C., Hilton-Taylor, C., Hoffmann, M., Isaac, N.J.B., Iversen, L.L., Outhwaite, C.L., Visconti, P., Geldmann, J., 2021. Using the IUCN Red List to map threats to terrestrial vertebrates at global scale. Nature Ecology & Evolution 5, 1510–1519.

IUCN, 2012. IUCN Red List Categories and Criteria. Version 3.1, Second edn, Gland, Switzerland and Cambridge, UK: IUCN. iv + 32pp.

IUCN, 2018. Mapping Standards and Data Quality for the IUCN Red List Categories and Criteria. Version 1.16.

IUCN, 2021. IUCN Red List of Threatened Species Version 2021-2

Jetz, W., Pyron, R.A., 2018. The interplay of past diversification and evolutionary isolation with present imperilment across the amphibian tree of life. Nature Ecology & Evolution 2, 850–858.

Karger, D.N., Conrad, O., Böhner, J., Kawohl, T., Kreft, H., Soria-Auza, R.W., Zimmermann, N.E., Linder, H.P., Kessler, M., 2017. Climatologies at high resolution for the earth’s land surface areas. Scientific data 4, 170122.

Karger, D.N., Conrad, O., Böhner, J., Kawohl, T., Kreft, H., Soria-Auza, R.W., Zimmermann, N.E., Linder, H.P., Kessler, M., 2018. Data from: Climatologies at high resolution for the earth’s land surface areas, Dryad, Dataset.

Lertzman-Lepofsky, G.F., Kissel, A.M., Sinervo, B., Palen, W.J., 2020. Water loss and temperature interact to compound amphibian vulnerability to climate change. Global Change Biology 26, 4868–4879.

Liedtke, H.C., 2019. AmphiNom: an amphibian systematics tool. Systematics and biodiversity 17, 1–6.

Lucas, P.M., González□Suárez, M., Revilla, E., 2019. Range area matters, and so does spatial configuration: predicting conservation status in vertebrates. Ecography.

Miller, D.A.W., Grant, E.H.C., Muths, E., Amburgey, S.M., Adams, M.J., Joseph, M.B., Waddle, J.H., Johnson, P.T.J., Ryan, M.E., Schmidt, B.R., Calhoun, D.L., Davis, C.L., Fisher, R.N., Green, D.M., Hossack, B.R., Rittenhouse, T.A.G., Walls, S.C., Bailey, L.L., Cruickshank, S.S., Fellers, G.M., Gorman, T.A., Haas, C.A., Hughson, W., Pilliod, D.S., Price, S.J., Ray, A.M., Sadinski, W., Saenz, D., Barichivich, W.J., Brand, A., Brehme, C.S., Dagit, R., Delaney, K.S., Glorioso, B.M., Kats, L.B., Kleeman, P.M., Pearl, C.A., Rochester, C.J., Riley, S.P.D., Roth, M., Sigafus, B.H., 2018. Quantifying climate sensitivity and climate-driven change in North American amphibian communities. Nature communications 9, 3926.

NASA, 2018. Gridded Population of the World, Version 4 (GPWv4): Population Density, Revision 11. NASA Socioeconomic Data and Applications Center (SEDAC), Palisades, NY.

Oliveira, B.F., São-Pedro, V.A., Santos-Barrera, G., Penone, C., Costa, G.C., 2017. AmphiBIO, a global database for amphibian ecological traits. Scientific data 4, 170123.

Olson, D.M., Dinerstein, E., Wikramanayake, E.D., Burgess, N.D., Powell, G.V.N., Underwood, E.C., D’amico, J.A., Itoua, I., Strand, H.E., Morrison, J.C., Loucks, C.J., Allnutt, T.F., Ricketts, T.H., Kura, Y., Lamoreux, J.F., Wettengel, W.W., Hedao, P., Kassem, K.R., 2001. Terrestrial Ecoregions of the World: A New Map of Life on Earth. Bioscience 51, 933–938, 936.

Penone, C., Davidson, A.D., Shoemaker, K.T., Di Marco, M., Rondinini, C., Brooks, T.M., Young, B.E., Graham, C.H., Costa, G.C., 2014. Imputation of missing data in life-history trait datasets: which approach performs the best? Methods in Ecology and Evolution 5, 961–970.

Pereira, H.M., Leadley, P.W., Proença, V., Alkemade, R., Scharlemann, J.P.W., Fernandez-Manjarrés, J.F., Araújo, M.B., Balvanera, P., Biggs, R., Cheung, W.W.L., Chini, L., Cooper, H.D., Gilman, E.L., Guénette, S., Hurtt, G.C., Huntington, H.P., Mace, G.M., Oberdorff, T., Revenga, C., Rodrigues, P., Scholes, R.J., Sumaila, U.R., Walpole, M., 2010. Scenarios for global biodiversity in the 21st century. Science 330, 1496–1501.

Pincheira-Donoso, D., Harvey, L.P., Cotter, S.C., Stark, G., Meiri, S., Hodgson, D.J., 2021a. The global macroecology of brood size in amphibians reveals a predisposition of low-fecundity species to extinction. Global Ecology and Biogeography 30, 1299–1310.

Pincheira-Donoso, D., Harvey, L.P., Grattarola, F., Jara, M., Cotter, S.C., Tregenza, T., Hodgson, D.J., 2021b. The multiple origins of sexual size dimorphism in global amphibians. Global Ecology and Biogeography 30, 443–458.

Ramalho, Q., Tourinho, L., Almeida□Gomes, M., Vale, M.M., Prevedello, J.A., 2021. Reforestation can compensate negative effects of climate change on amphibians. Biological Conservation 260, 109187.

Roberts, D.R., Bahn, V., Ciuti, S., Boyce, M.S., Elith, J., Guillera-Arroita, G., Hauenstein, S., Lahoz-Monfort, J.J., Schröder, B., Thuiller, W., Warton, D.I., Wintle, B.A., Hartig, F., Dormann, C.F., 2017. Cross-validation strategies for data with temporal, spatial, hierarchical, or phylogenetic structure. Ecography 40, 913–929.

Rondinini, C., Di Marco, M., Visconti, P., Butchart, S.H.M., Boitani, L., 2014. Update or Outdate: Long-Term Viability of the IUCN Red List. Conservation Letters 7, 126–130.

Scheele, B.C., Pasmans, F., Skerratt, L.F., Berger, L., Martel, A., Beukema, W., Acevedo, A.A., Burrowes, P.A., Carvalho, T., Catenazzi, A., De la Riva, I., Fisher, M.C., Flechas, S.V., Foster, C.N., Frías-Álvarez, P., Garner, T.W.J., Gratwicke, B., Guayasamin, J.M., Hirschfeld, M., Kolby, J.E., Kosch, T.A., La Marca, E., Lindenmayer, D.B., Lips, K.R., Longo, A.V., Maneyro, R., McDonald, C.A., Mendelson, J., Palacios-Rodriguez, P., Parra-Olea, G., Richards-Zawacki, C.L., Rödel, M.-O., Rovito, S.M., Soto-Azat, C., Toledo, L.F., Voyles, J., Weldon, C., Whitfield, S.M., Wilkinson, M., Zamudio, K.R., Canessa, S., 2019. Amphibian fungal panzootic causes catastrophic and ongoing loss of biodiversity. Science 363, 1459–1463.

Sodhi, N.S., Bickford, D., Diesmos, A.C., Lee, T.M., Koh, L.P., Brook, B.W., Sekercioglu, C.H., Bradshaw, C.J.A., 2008. Measuring the Meltdown: Drivers of Global Amphibian Extinction and Decline. PLOS ONE 3, e1636.

Stekhoven, D.J., Bühlmann, P., 2011. MissForest—non-parametric missing value imputation for mixed-type data. Bioinformatics 28, 112–118.

Stuart, S.N., Chanson, J.S., Cox, N.A., Young, B.E., Rodrigues, A.S.L., Fischman, D.L., Waller, R.W., 2004. Status and Trends of Amphibian Declines and Extinctions Worldwide. Science 306, 1783–1786.

Stuart, S.N., Hoffmann, M., Chanson, J.S., Cox, N.A., Berridge, R.J., Ramani, P., Young, B.E., 2008. Threatened Amphibians of the World. Lynx Edicions, Barcelona, Spain; IUCN, Gland, Switzerland; and Conservation International, Arlington, Virginia, USA.

Tange, R.I., Rasmussen, M.A., Taira, E., Bro, R., 2017. Benchmarking support vector regression against partial least squares regression and artificial neural network: Effect of sample size on model performance. Journal of Near Infrared Spectroscopy 25, 381–390.

Tapley, B., Michaels, C.J., Gumbs, R., Böhm, M., Luedtke, J., Pearce-Kelly, P., Rowley, J. J.L., 2018. The disparity between species description and conservation assessment: A case study in taxa with high rates of species discovery. Biological Conservation 220, 209–214.

Urban, M.C., Bocedi, G., Hendry, A.P., Mihoub, J.B., Peer, G., Singer, A., Bridle, J.R., Crozier, L.G., Meester, D.L., Godsoe, W., Gonzalez, A., Hellmann, J.J., Holt, R.D., Huth, A., Johst, K., Krug, C.B., Leadley, P.W., Palmer, S.C.F., Pantel, J.H., Schmitz, A., Zollner, P.A., Travis, J.M.J., 2016. Improving the forecast for biodiversity under climate change. Science 353.

Vabalas, A., Gowen, E., Poliakoff, E., Casson, A.J., 2019. Machine learning algorithm validation with a limited sample size. PLOS ONE 14, e0224365.

Wake, D.B., Vredenburg, V.T., 2008. Are we in the midst of the sixth mass extinction? A view from the world of amphibians. Proceedings of the National Academy of Sciences 105, 11466–11473.

Weiss, D.J., Nelson, A., Gibson, H.S., Temperley, W., Peedell, S., Lieber, A., Hancher, M., Poyart, E., Belchior, S., Fullman, N., Mappin, B., Dalrymple, U., Rozier, J., Lucas, T.C.D., Howes, R.E., Tusting, L.S., Kang, S.Y., Cameron, E., Bisanzio, D., Battle, K.E., Bhatt, S., Gething, P.W., 2018. A global map of travel time to cities to assess inequalities in accessibility in 2015. Nature 553, 333–336.

Wieringa, J.G., 2022. Comparing predictions of IUCN Red List categories from machine learning and other methods for bats. Journal of Mammalogy 103, 528–539.

Zizka, A., Andermann, T., Silvestro, D., 2022. IUCNN - Deep learning approaches to approximate species’ extinction risk. Diversity and Distributions 28, 227–241.

Zizka, A., Silvestro, D., Vitt, P., Knight, T.M., 2021. Automated conservation assessment of the orchid family with deep learning. Conservation Biology 35, 897–908.

