## Supplementary material for "Testing the predictive performance of comparative extinction risk models to support the global amphibian assessment": Table A1

**Table A1.** Description and source of the variables used for the imputation of species traits.

| **Variable Group** | **Description** | **Source** |
| --- | --- | --- |
| ***Variable*** |  |  |
| Taxonomy |  |  |
| *Order* | Factorial variable with three levels indicating the taxonomic order of the species (*Anura*, *Caudata* or *Gymnophiona*) | IUCN 2021 |
| Spatial context |  |  |
| *Realm* | Factorial variable indicating the most representative realm where the species lives, with a numerical code ranging from 1 to 8 representing *Australasian*, *Antarctica*, *Afrotropical*, *Indomalayan*, *Neartic*, *Neotropical*, *Oceanian* and *Paleartic* respectively | Olson et al. 2004 |
| *Realm_Australasian* | If the species lives or not in this realm | Olson et al. 2004 |
| *Realm_Antartica* | If the species lives or not in this realm | Olson et al. 2004 |
| *Realm_Afrotropical* | If the species lives or not in this realm | Olson et al. 2004 |
| *Realm_Indomalayan* | If the species lives or not in this realm | Olson et al. 2004 |
| *Realm_Neartic* | If the species lives or not in this realm | Olson et al. 2004 |
| *Realm_Neotropical* | If the species lives or not in this realm | Olson et al. 2004 |
| *Realm_Oceanian* | If the species lives or not in this realm | Olson et al. 2004 |
| *Realm_Paleartic* | If the species lives or not in this realm | Olson et al. 2004 |
| *N_Realm* | Number of realms where the species lives | Olson et al. 2004 |
| Climate |  |  |
| *Bio_1_mean* | Mean of the respective bioclimatic variable within the species range | Karger et al. 2017 |
| *Bio_2_mean* | Mean of the respective bioclimatic variable within the species range | Karger et al. 2017 |
| *Bio_3_mean* | Mean of the respective bioclimatic variable within the species range | Karger et al. 2017 |
| *Bio_4_mean* | Mean of the respective bioclimatic variable within the species range | Karger et al. 2017 |
| *Bio_5_mean* | Mean of the respective bioclimatic variable within the species range | Karger et al. 2017 |
| *Bio_6_mean* | Mean of the respective bioclimatic variable within the species range | Karger et al. 2017 |
| *Bio_7_mean* | Mean of the respective bioclimatic variable within the species range | Karger et al. 2017 |
| *Bio_8_mean* | Mean of the respective bioclimatic variable within the species range | Karger et al. 2017 |
| *Bio_9_mean* | Mean of the respective bioclimatic variable within the species range | Karger et al. 2017 |
| *Bio_10_mean* | Mean of the respective bioclimatic variable within the species range | Karger et al. 2017 |
| *Bio_11_mean* | Mean of the respective bioclimatic variable within the species range | Karger et al. 2017 |
| *Bio_12_mean* | Mean of the respective bioclimatic variable within the species range | Karger et al. 2017 |
| *Bio_13_mean* | Mean of the respective bioclimatic variable within the species range | Karger et al. 2017 |
| *Bio_14_mean* | Mean of the respective bioclimatic variable within the species range | Karger et al. 2017 |
| *Bio_15_mean* | Mean of the respective bioclimatic variable within the species range | Karger et al. 2017 |
| *Bio_16_mean* | Mean of the respective bioclimatic variable within the species range | Karger et al. 2017 |
| *Bio_17_mean* | Mean of the respective bioclimatic variable within the species range | Karger et al. 2017 |
| *Bio_18_mean* | Mean of the respective bioclimatic variable within the species range | Karger et al. 2017 |
| *Bio_19_mean* | Mean of the respective bioclimatic variable within the species range | Karger et al. 2017 |
| *Bio_1_sd* | SD of the respective bioclimatic variable within the species range | Karger et al. 2017 |
| *Bio_2_sd* | SD of the respective bioclimatic variable within the species range | Karger et al. 2017 |
| *Bio_3_sd* | SD of the respective bioclimatic variable within the species range | Karger et al. 2017 |
| *Bio_4_sd* | SD of the respective bioclimatic variable within the species range | Karger et al. 2017 |
| *Bio_5_sd* | SD of the respective bioclimatic variable within the species range | Karger et al. 2017 |
| *Bio_6_sd* | SD of the respective bioclimatic variable within the species range | Karger et al. 2017 |
| *Bio_7_sd* | SD of the respective bioclimatic variable within the species range | Karger et al. 2017 |
| *Bio_8_sd* | SD of the respective bioclimatic variable within the species range | Karger et al. 2017 |
| *Bio_9_sd* | SD of the respective bioclimatic variable within the species range | Karger et al. 2017 |
| *Bio_10_sd* | SD of the respective bioclimatic variable within the species range | Karger et al. 2017 |
| *Bio_11_sd* | SD of the respective bioclimatic variable within the species range | Karger et al. 2017 |
| *Bio_12_sd* | SD of the respective bioclimatic variable within the species range | Karger et al. 2017 |
| *Bio_13_sd* | SD of the respective bioclimatic variable within the species range | Karger et al. 2017 |
| *Bio_14_sd* | SD of the respective bioclimatic variable within the species range | Karger et al. 2017 |
| *Bio_15_sd* | SD of the respective bioclimatic variable within the species range | Karger et al. 2017 |
| *Bio_16_sd* | SD of the respective bioclimatic variable within the species range | Karger et al. 2017 |
| *Bio_17_sd* | SD of the respective bioclimatic variable within the species range | Karger et al. 2017 |
| *Bio_18_sd* | SD of the respective bioclimatic variable within the species range | Karger et al. 2017 |
| *Bio_19_sd* | SD of the respective bioclimatic variable within the species range | Karger et al. 2017 |
| Ecological traits |  |  |
| *Nesting_site* | Factorial variable with five levels describing where parents lay their eggs: *water*, *ground*, *burrows*, *vegetation*, *brooder* | Pincheira-Donoso et al. 2021a and Pincheira-Donoso et al. 2021b |
| *Body_size* | For Anura and Caudata, we used maximum snout–vent length (SVL) as our measure of body size, given that this is the most widely used proxy for body size in these two orders. For Gymnophiona, we used the maximum total body length (Pincheira-Donoso et al., 2019). All these values were log transformed | Pincheira-Donoso et al. 2021a |
| *Body_size_Males* | Body size variable for males | Pincheira-Donoso et al. 2021a |
| *Body_size_Females* | Body size variable for females | Pincheira-Donoso et al. 2021a |
| *SSD* | Species sexual size dimorphism (SSD) | Pincheira-Donoso et al. 2021b |
| *Brood_size* | The number of offspring or eggs per clutch log transformed | Pincheira-Donoso et al. 2021a and Oliveira et al. 2017 |
| *Brood_size_min* | The minimum number of offspring or eggs per clutch log transformed | Pincheira-Donoso et al. 2021a and Oliveira et al. 2017 |
| *Brood_size_max* | The maximum number of offspring or eggs per clutch log transformed | Pincheira-Donoso et al. 2021a and Oliveira et al. 2017 |
| *Offspring_size_min* | Minimum offspring or egg size (millimeter) log transformed | Oliveira et al. 2017 |
| *Offspring_size_max* | Maximum offspring or egg size (millimeter) log transformed | Oliveira et al. 2017 |
| *Reproductive_output* | Maximum number reproduction events per year log transformed | Oliveira et al. 2017 |
| *Parity* | Factorial variable with three levels indicating how the species reproduces: *direct*, *larval* or *viviparous*. | Oliveira et al. 2017 |
| *Habitat_generalist_1* | Whether the species is generalist (is in more than one habitat) or not | IUCN 2021 |
| *Habitat_generalist_2* | Whether the species is generalist (is in more than two habitats) or not | IUCN 2021 |
| *N_Habitat* | Number of habitats where the species lives | IUCN 2021 |
| *Microhabitat_Fos* | Species lives in fossorial habitat | Oliveira et al. 2017 |
| *Microhabitat_Ter* | Species lives in terrestrial habitat | Oliveira et al. 2017 |
| *Microhabitat_Aqu* | Species lives in aquatic habitat | Oliveira et al. 2017 |
| *Microhabitat_Arb* | Species lives in arboreal habitat | Oliveira et al. 2017 |
| *Activity_Diurnal* | The species is active during the day | Oliveira et al. 2017 |
| *Activity_Crepuscular* | Crepuscular (i.e., active during the period immediately after dawn and that immediately before dusk) | Oliveira et al. 2017 |
| *Activity_Nocturnal* | The species is active during the night | Oliveira et al. 2017 |
| *Seas_act_Wet_warm* | Describe if the species is active is during wet and warm months. Seasonal period as active. Based on the comparison of the precipitation (wet or dry) and temperature (warm or cold) conditions when active in relation to the average climatic conditions over the year. Climatic conditions were obtained from weather stations closer to localities where specimens were collected or to field sites reported in publications (available at www.weatherbase.com). | Oliveira et al. 2017 |
| *Seas_act_Wet_cold* | Describe if the species is active during wet and cold months. Seasonal period as active. Based on the comparison of the precipitation (wet or dry) and temperature (warm or cold) conditions when active in relation to the average climatic conditions over the year. Climatic conditions were obtained from weather stations closer to localities where specimens were collected or to field sites reported in publications (available at www.weatherbase.com). | Oliveira et al. 2017 |
| *Sea_acts_Dry_warm* | Describe if the species is active during dry and warm months. Seasonal period as active. Based on the comparison of the precipitation (wet or dry) and temperature (warm or cold) conditions when active in relation to the average climatic conditions over the year. Climatic conditions were obtained from weather stations closer to localities where specimens were collected or to field sites reported in publications (available at www.weatherbase.com). | Oliveira et al. 2017 |
| *Sea_acts_Dry_cold* | Describe if the species is active during dry and cold months. Seasonal period as active. Based on the comparison of the precipitation (wet or dry) and temperature (warm or cold) conditions when active in relation to the average climatic conditions over the year. Climatic conditions were obtained from weather stations closer to localities where specimens were collected or to field sites reported in publications (available at www.weatherbase.com). | Oliveira et al. 2017 |
| *Diet_Flowers* | Species eat flowers. Food items from the eating habits of adults using qualitative dietary categories. Information is based of specialist guess, direct observation or stomach content examination, as reported in the literature | Oliveira et al. 2017 |
| *Diet_Seeds* | Species eat seeds. Food items from the eating habits of adults using qualitative dietary categories. Information is based of specialist guess, direct observation or stomach content examination, as reported in the literature | Oliveira et al. 2017 |
| *Diet_Fruits* | Species eat fruits. Food items from the eating habits of adults using qualitative dietary categories. Information is based of specialist guess, direct observation or stomach content examination, as reported in the literature | Oliveira et al. 2017 |
| *Diet_Artrophods* | Species eat arthropods. Food items from the eating habits of adults using qualitative dietary categories. Information is based of specialist guess, direct observation or stomach content examination, as reported in the literature | Oliveira et al. 2017 |
| *Diet_Vertebrates* | Species eat vertebrates. Food items from the eating habits of adults using qualitative dietary categories. Information is based of specialist guess, direct observation or stomach content examination, as reported in the literature | Oliveira et al. 2017 |
| *Diet_Leaves* | Species eat leaves. Food items from the eating habits of adults using qualitative dietary categories. Information is based of specialist guess, direct observation or stomach content examination, as reported in the literature | Oliveira et al. 2017 |
| Range/Spatial configuration |  |  |
| *Range_Area_original* | Area in square kilometers of the species range log transformed | IUCN 2021 |
| *Phylogeny* |  |  |
| *Phylo_PC1* | First principal component from phylogeny | Jetz et al. 2019 |
| *Phylo_PC2* | Second principal component from phylogeny | Jetz et al. 2019 |
| *Phylo_PC3* | Third principal component from phylogeny | Jetz et al. 2019 |
| *Phylo_PC4* | Fourth principal component from phylogeny | Jetz et al. 2019 |
| *Phylo_PC5* | Fifth Principal component from phylogeny | Jetz et al. 2019 |
| *Phylo_PC6* | Sixth principal component from phylogeny | Jetz et al. 2019 |

**Supplementary methods**

*Climate change variable calculation*

We quantified recent climate change within species distribution, using CHELSA monthly time series (Karger et al. 2017, 2018). We calculated the average climatic conditions for two periods of time: (1) between January 1965 and December 1994, *Historical climate*, and (2) between January 2005 and December 2014, *Current climate*. We then used the ‘dismo’ package (Hijmans et al. 2020), to calculate the 19 bioclimatic variables for each period of time. Using the rasterized distributions (See section *Species distributions and spatial variables* within the Material and Methods section of the main text), we calculated the mean values for each bioclimatic variable and period for each species. Then, we merged the mean values for each bioclimatic variable per species and for each period, which represented the bioclimatic space for historical and current for amphibians. Over these data we run a principal component analysis using function *prcomp* in R and selected the five first components which explained 93% of the variance (Table A6). We then used the two values, one for each period, of each component for each species and calculated the change in the value of the component as the difference of the value of the component in the *Current climate* minus the value of the component in the *Historical climate*, obtaining five variables describing climate change: *Climate_Change_1*, *Climate_Change_2*, *Climate_Change_3*, *Climate_Change_4* and *Climate_Change_5* representing the changes in the first, second, third, fourth and fifth principal components respectively.

*Descriptors of the geographic distribution range*

We projected the selected range polygons from the IUCN Red List dataset (IUCN 2021; see Species distributions and spatial variables on Material and methods section in the main text) to Winkel tripel (WT) projection instead of an equal area, to minimize errors in shapes, distances, and areas. Using the data of length and area we calculated *Range_Circularity*, a descriptor for the fragment shape.

*Range_Circularity* = *P_H_*/*P_O_*

Where *P_H_* = total perimeter of idealized fragments with the same area as the observed fragments but with circular shape, and *P_O_* = actual total perimeter of the fragments, and it ranges from 0 (most irregular shapes) to 1 (completely circular shapes; Lucas et al. 2019).

*Variables and databases modifications for models of extinction risk*

To reduce the number of bioclimatic variables in our models, we applied a principal component analysis (PCA). We calculated the PCA for the values of the mean and the standard deviation of the 19 bioclimatic variables measured within each species range, then selected the first four components that explained the 74% of the variance (Table A5).

Habitat variables used for data imputation (Table A1) were modified for the models of extinction risk to have more balanced groups. We created a new factorial variable with four levels called *Hab_ generalist _forest* indicating whether the species is generalist (lives in more than 1 habitat) and is or not in forest habitat: *Generalist_Forest*, *Generalist_NoForest*, *Specialist_Forest*, *Specialist_NoForest* (Table A7). Shrubland and grassland species were merged because 0.7% only of analyzed amphibians were associated exclusively to shrublands while, if generalists were excluded, 81% of shrubland amphibians were also associated to grasslands. Several studies have shown that, in amphibian communities, a gradient exist between forest specialists and species living in more open and sunny habitats (Ficetola et al. 2015; Nowakowski et al. 2017; Skelly et al. 1999).

Because we used different model algorithms to calculate extinction risk (see below), which require different formats for the predictor variables (numeric or factorial, or only numeric), we generated two different datasets. One dataset including numeric and factorial variables, the second, including numeric and boolean variables, which were converted from the factorial variables (e.g. from *Parity* factorial variable with “*Direct*”, “*Larvae*” and ‘*Viviparous’* levels, to “*Parity_Direct”* and “*Parity_Larvae”* and “*Parity_Viviparous”* as distinct boolean variables). We converted to boolean variables the factorial variables *Realm*, *Parity*, and *Generalist* (Table A7)*.*

In addition, the four original boolean variables (*Microhab_Fos, Microhab_Ter, Microhab_Aqu, Microhab_Arb*) describing if the species lives (1) or not (0) in a microhabitat, were aggregated into a unique factor variable. Such as these four boolean variables are not exclusive, e.g. a species can be present in *Microhab_Fos* and in *Microhab_Arb*, that means that there are multiple possible combinations of these four boolean variables. In total we observed 14 realized combinations. Due to inclusion of a factor variable with so many levels and only few species in some of these levels was causing problems to fit and validate the models, we regrouped de variables to three levels (*Semiaquatic*, *Generalist* and *Others*) to have a more balanced variable with respect to the number species while describing the generalist/specialist characteristics at microhabitat level which may influence the risk of extinction of the species (Table A8).

In order to avoid an over precision / exactitude of the area of the range, we modified the original value truncating the *Range_area_original*.

*Range_area = truncation (Range_area_original)*

Some variables were manually excluded when some model algorithms and/or the validation process of the respective models were not possible with such variables. This happened for binary and/or factorial variables which were highly unbalance such as *Microhab_Fos* for Gymnophiona with all species occupying this microhabitat (See Table A11).

*Modelling extinction risk*

For cumulative link models (CLM) we used the databases with numeric and boolean variables. Using the non-correlated variables, we first fitted a full model with all variables using the clm function from the package ordinal (Christensen 2020) applying a probit link function. To account with unbalanced distribution among categories, we used weights proportional to the number of species in each category. Over this full model we performed a stepwise backward model selection procedure by AIC using ‘MASS’ package (Venables and Ripley 2002).

For phylogenetic generalised least squares models (PGLS) we used the databases with factorial variables and the four phylogenetic eigen vectors were removed from the database. With all the non-correlated variables, using gls function in package nlme (Pinheiro et al. 2021), we first fitted a full GLS model, which included the phylogeny, using the ‘corBrownian’ function from package ‘ape’ (Paradis and Schliep 2019), as a correlation matrix among the phylogenetic distances. As with the CLMs we performed a stepwise backward model selection procedure by AIC in MASS package (Venables and Ripley 2002). Predicted values were reclassified in the five Red List levels: LC for < 1.5, NT for 1.5 - 2.5, VU for 2.5 - 3.5, EN for 3.5-4.5 and CR for > 4.5.

For random forest (RF) models we used the databases with factorial variables and as response variable (Red List categories) as factor. We used randomForest package (Liaw and Wiener 2002) to fit the models. Although RF models can operate with large numbers of variables, this can lead to an increase in the correlation of trees, reducing the overall performance of the model. To avoid this problem, using all the non-correlated variables, we applied a RF model selection approach using function rf.modelSel from package rfUtilities (Evans and Murphy 2018) which apply the procedure described in Murphy et al 2010. We used "mir" option for scaling importance values, a vector 100 percentiles values to test (r), a mtry value equals to the square root of the number of variables and 2,000 trees.

For neural network models (NN) we used the databases with numeric and boolean variables and as response variable a numeric variable with the categories of the IUCN (regression NN). We used h2o package to fit the models (LeDell et al. 2022) in R. NN models have many parameters and hyperparameters to adjust. We applied a process of hyper-parameter optimization for multilayer artificial neural network models considering 648 different potential models which resulted from combinations of three different activation parameters ("Tanh", "RectifierWithDropout", "TanhWithDropout"), four options for hidden layers ([349, 174, 87, 29]; [174, 87, 29]; [87, 29]; [27, 9]), six input-dropout ratio options (0.05, 0.1, 0.15, 0.2, 0.3, 0.4), three options for Lasso regularization (10^-3^, 10^-4^, 10^-5^) and three options for Ridge regularization (10^-3^, 10^-4^, 10^-5^). The search criteria within the grid of the potential models was using a random discrete strategy.

**Table A2.** Variables preselected for comparative analysis of extinction risk. We add the description, source and the reference of the justification for the inclusion of each variable.

| **Variable Group** | **Description** | **Source** | **Justification** |
| --- | --- | --- | --- |
| ***Variable*** |  |  |  |
| **Land cover/Land cover change** |  |  |  |
| *Urbanization* | Percentage of urban areas in the species range | ESA 2021 | McKinney 2002; Newbold et al. 2015 |
| *Agriculture* | Percentage of agriculture areas in the species range | ESA 2021 | Newbold et al. 2015 |
| *Urbanization_Change* | Change in *Urban* in 10 years (*Urban_t_0_* – *Urban_t_1_*) | ESA 2021 | Newbold et al. 2015 |
| *Agriculture_Change* | Change in *Agriculture* in 10 years (*Agriculture_t_0_* – *Agriculture_t_1_*) | ESA 2021 | Newbold et al. 2015 |
| *Human_density* | Human population density (humans/km^2^) | NASA 2018 | Newbold et al. 2015 |
| *Human_density_Change* | Change in *Human* in 10 years (*Human_t_0_* – *Human_t_1_*) |  | Newbold et al. 2015 |
| *Accessibility* | Travel time to cities (higher values correspond to more inaccessible areas) | Weiss et al. 2018 | Benítez-López et al. 2019 |
| **Spatial context** |  |  |  |
| *Realm* | The most representative realm where the species lives, with a numerical code ranging from 1 to 8 representing Australasian, Antarctica, Afrotropical, Indomalayan, Neartic, Neotropical, Oceanian and Paleartic respectively | Olson et al. 2001 | Yackulic et al 2011 |
| **Climate** |  |  |  |
| *Climate_1* | First principal component from climate | Karger et al. 2017, 2018 | Hof et al. 2011; Silvano & Segalla 2005 |
| *Climate_2* | Second principal component from climate | Karger et al. 2017, 2018 | Hof et al. 2011; Silvano & Segalla 2005 |
| *Climate_3* | Third principal component from climate | Karger et al. 2017, 2018 | Hof et al. 2011; Silvano & Segalla 2005 |
| *Climate_4* | Fourth principal component from climate | Karger et al. 2017, 2018 | Hof et al. 2011; Silvano & Segalla 2005 |
| **Climate Change** |  |  |  |
| *Climate_Change_1* | First principal component from climate change | Karger et al. 2017, 2018 | Urban 2015; Hof et al. 2011; Silvano & Segalla 2005 |
| *Climate_Change_2* | Second principal component from climate change | Karger et al. 2017, 2018 | Urban 2015; Hof et al. 2011; Silvano & Segalla 2005 |
| *Climate_Change_3* | Third principal component from climate change | Karger et al. 2017, 2018 | Urban 2015; Hof et al. 2011; Silvano & Segalla 2005 |
| *Climate_Change_4* | Fourth principal component from climate change | Karger et al. 2017, 2018 | Urban 2015; Hof et al. 2011; Silvano & Segalla 2005 |
| *Climate_Change_5* | Fifth principal component from climate change | Karger et al. 2017, 2018 | Urban 2015; Hof et al. 2011; Silvano & Segalla 2005 |
| **Ecological traits** |  |  |  |
| *Body_size* | For anurans and salamanders, we used maximum snout–vent length (SVL) as our measure of body size, given that this is the most widely used proxy for body size in these two orders. For caecilians, maximum total body length is the traditional measure of size (Pincheira-Donoso et al. 2019), and thus the proxy we used. All were log transformed | Pincheira-Donoso et al. 2021 | Cardillo et al. 2008; Cardillo 2021 |
| *Brood_size* | The brood size of the species log transformed | Pincheira-Donoso et al. 2021 and Oliveira et al. 2017 | Pincheira-Donoso et al. 2021 |
| *Parity* | Factor variable indicating whether the species reproduce via direct, larval development or is viviparous. | Oliveira et al. 2017 |  |
| *Habitat_generalist* | Wheter the species is generalist (is in more than one habitat) or not | IUCN 2021 | Carilo Filho et al. 2021 |
| *Hab_ generalist _forest* | Factorial variable with four levels indicating whether the species is generalist (lives in more than 1 habitat) and is or not in forest habitat: *Generalist_Forest*, *Generalist_NoForest*, *Specialist_Forest*, *Specialist_NoForest* | IUCN 2021 | Carilo Filho et al. 2021 |
| *Microhabitat_Fos* | Species lives in fossorial habitat | Oliveira et al. 2017 | Carilo Filho et al. 2021 |
| *Microhabitat_Ter* | Species lives in terrestrial habitat | Oliveira et al. 2017 | Carilo Filho et al. 2021 |
| *Microhabitat_Aqu* | Species lives in aquatic habitat | Oliveira et al. 2017 | Carilo Filho et al. 2021 |
| *Microhabitat_Arb* | Species lives in arboreal habitat | Oliveira et al. 2017 | Carilo Filho et al. 2021 |
| **Range/Spatial configuration** |  |  |  |
| *Range_area* | Area in square kilometers of the species range log transformed and then truncated | IUCN 2021 | Lucas et al. 2019 |
| *Range_fragments* | The number of fragments in which is divided the species range | IUCN 2021 | Lucas et al. 2019 |
| *Range_Circularity* | Shape ratio calculated as PH/PO, where PH = total perimeter of idealized fragments with the same area as the observed fragments but with circular shape, andPO = actual total perimeter of the fragments. The ratio ranges from 0 (most irregular shapes) to 1 (completely circular shapes). | IUCN 2021 | Lucas et al. 2019 |
| *Range_heterogeinity* | Proportion of the total range area represented by the largest fragment. It ranges from close to 0 (similar fragment size) to close to 1 (very different fragment size). | IUCN 2021 | Lucas et al. 2019 |
| **Phylogeny** |  |  |  |
| *Phylogeny_1* | First principal component from phylogeny | Jetz et al. 2019 | González-del-Pliego et al. 2019 |
| *Phylogeny_2* | Second principal component from phylogeny | Jetz et al. 2019 | González-del-Pliego et al. 2019 |
| *Phylogeny_3* | Third principal component from phylogeny | Jetz et al. 2019 | González-del-Pliego et al. 2019 |
| *Phylogeny_4* | Fourth principal component from phylogeny | Jetz et al. 2019 | González-del-Pliego et al. 2019 |

**Table A3.** Results of principal component analysis (PCA) for phylogenetic tree for amphibians (Jetz et al. 2019) showing the first 10 components. For data imputation we chosen the first six principal components, which explained the 96% of the variance. For modelling extinction risk, in order to reduce the number of variables in the models, we selected the first four principal components which explained the 93% of the variance.

|  | **PC1** | **PC2** | **PC3** | **PC4** | **PC5** | **PC6** | **PC7** | **PC8** | **PC9** | **PC10** |
| --- | --- | --- | --- | --- | --- | --- | --- | --- | --- | --- |
| **Proportion of variance explained** | 0.56 | 0.31 | 0.04 | 0.02 | 0.02 | 0.01 | 0.01 | 0.01 | 0.00 | 0.00 |
| **Cumulative proportion of variance explained** | 0.56 | 0.87 | 0.90 | 0.93 | 0.94 | 0.96 | 0.96 | 0.97 | 0.97 | 0.98 |

**Table A4.** Results for the data imputation analysis for species traits. We show the Normalized Root Mean Square Error (NRMSE) and the proportion of missing data (Missingness) for each species trait variable.

| **Variable** | **NRMSE** | **Missingness** |
| --- | --- | --- |
| *N _realm* | 0.43 | 0.04 |
| *N _habitat_lvl_0* | 0.59 | 0.00 |
| *Range_area_original* | 0.34 | 0.03 |
| *Body_size* | 0.26 | 0.23 |
| *SVL_Males* | 0.27 | 0.56 |
| *SVL_Females* | 0.25 | 0.56 |
| *SSD* | 0.77 | 0.56 |
| *Brood_size* | 0.25 | 0.72 |
| *Brood_size_min* | 0.36 | 0.77 |
| *Brood_size_max* | 0.31 | 0.77 |
| *Offspring_size_min* | 0.50 | 0.81 |
| *Offspring_size_max* | 0.48 | 0.81 |
| *Reproductive_output* | 0.98 | 0.39 |
| *Phylogeny_1* | 0.03 | 0.07 |
| *Phylogeny_2* | 0.04 | 0.07 |
| *Phylogeny_3* | 0.05 | 0.07 |
| *Phylogeny_4* | 0.04 | 0.07 |
| *Phylogeny_5* | 0.04 | 0.07 |
| *Phylogeny_6* | 0.07 | 0.07 |

**Figure A1.** Results for the data imputation analysis for species traits. We show the Normalized Root Mean Square Error (NRMSE) and the proportion of missing data (Missingness) for each species trait variable. Vertical lines at 0.4 indicates the value to filter variables from imputation, we discarded trait variables with a NRMSE > 0.4 and a proportion of missing data > 0.4.

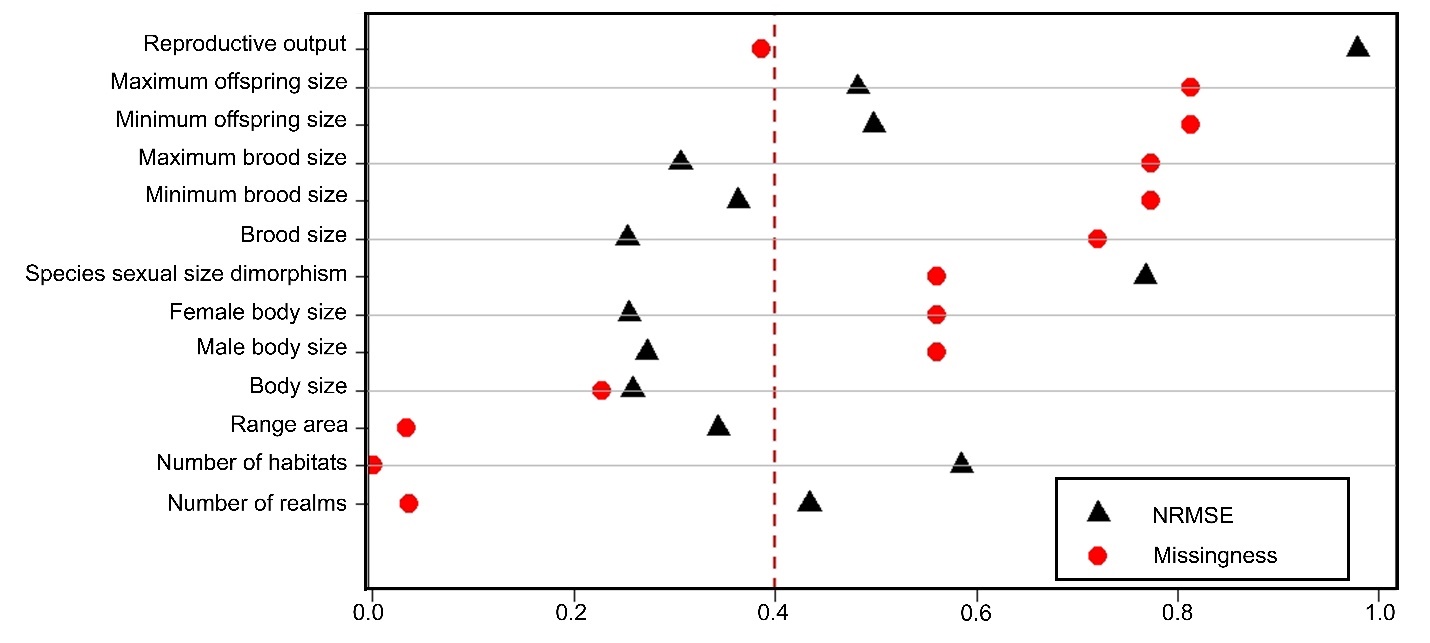

**Table A5.** Results of principal component analysis for bioclimatic variables used in the imputation showing the first 19 components. The four first components, which explain the 74% of the variance, were chosen for modelling extinction risk of species.

|  | **PC1** | **PC2** | **PC3** | **PC4** | **PC5** | **PC6** | **PC7** | **PC8** | **PC9** | **PC10** | **PC11** | **PC12** | **PC13** | **PC14** | **PC15** | **PC16** | **PC17** | **PC18** | **PC19** |
| --- | --- | --- | --- | --- | --- | --- | --- | --- | --- | --- | --- | --- | --- | --- | --- | --- | --- | --- | --- |
| **Proportion of variance explained** | 0.28 | 0.24 | 0.15 | 0.07 | 0.06 | 0.04 | 0.03 | 0.03 | 0.02 | 0.01 | 0.01 | 0.01 | 0.01 | 0.01 | 0.01 | 0.01 | 0.00 | 0.00 | 0.00 |
| **Cumulative proportion of variance explained** | 0.28 | 0.52 | 0.67 | 0.74 | 0.80 | 0.84 | 0.88 | 0.90 | 0.92 | 0.93 | 0.94 | 0.95 | 0.96 | 0.97 | 0.97 | 0.98 | 0.98 | 0.99 | 0.99 |

**Table A6.** Results of principal component analysis for climate change bioclimatic variables. The five first components, which explain the 93%, were chosen for modelling extinction risk of species.

|  | **PC1** | **PC2** | **PC3** | **PC4** | **PC5** | **PC6** | **PC7** | **PC8** | **PC9** | **PC10** | **PC11** | **PC12** | **PC13** | **PC14** | **PC15** | **PC16** | **PC17** | **PC18** | **PC19** |
| --- | --- | --- | --- | --- | --- | --- | --- | --- | --- | --- | --- | --- | --- | --- | --- | --- | --- | --- | --- |
| **Proportion of variance explained** | 0.44 | 0.23 | 0.12 | 0.09 | 0.05 | 0.03 | 0.01 | 0.01 | 0.01 | 0.00 | 0.00 | 0.00 | 0.00 | 0.00 | 0.00 | 0.00 | 0.00 | 0.00 | 0.00 |
| **Cumulative proportion of variance explained** | 0.44 | 0.67 | 0.79 | 0.88 | 0.93 | 0.97 | 0.98 | 0.99 | 0.99 | 1.00 | 1.00 | 1.00 | 1.00 | 1.00 | 1.00 | 1.00 | 1.00 | 1.00 | 1.00 |

**Table A7**. Conversions from factor variables to boolean variables.

| **Factor variable** | | **Boolean variables** | |
| --- | --- | --- | --- |
| **Name** | **Level** | **Name** | **Description** |
| *Realm* | *Australasian* | *Realm_Australasian* | If the species is mostly present in the Australasian realm (1) or not (0) |
|  | *Afrotropical* | *Realm_Afrotropical* | If the species is mostly present in the Afrotropical realm (1) or not (0) |
|  | *Indomalayan* | *Realm_Indomalayan* | If the species is mostly present in the Indomalayan realm (1) or not (0) |
|  | *Neartic* | *Realm_Neartic* | If the species is mostly present in the Neartic realm (1) or not (0) |
|  | *Neotropical* | *Realm_Neotropical* | If the species is mostly present in the Neotropical realm (1) or not (0) |
|  | *Paleartic* | *Realm_Paleartic* | If the species is mostly present in the Paleartic realm (1) or not (0) |
| *Parity* | *Direct* | *Parity_Direct* | Whether the species reproduce via direct development (1) or not (0) |
|  | *Larvae* | *Parity_Larvae* | Whether the species reproduce via larvae development (1) or not (0) |
|  | *Viviparous* | *Parity_Viviparous* | Whether the species reproduce via viviparous development (1) or not (0) |
| *Generalist* | *Generalist_NoForest* | *Generalist_NoForest* | Wheter the species is generalist (lives in more than 1 habitat) and is not in forest habitat (1) or not (0) |
|  | *Generalist_Forest* | *Generalist_Forest* | Wheter the species is generalist (lives in more than 1 habitat) and is in forest habitat (1) or not (0) |
|  | *Specialist_NoForest* | *Specialist_NoForest* | Wheter the species is specialist (lives in 1 habitat) and is not in forest habitat (1) or not (0) |
|  | *Specialist_Forest* | *Specialist_Forest* | Wheter the species is specialist (lives in 1 habitat) and is in forest habitat (1) or not (0) |

**Table A8**. Conversions from boolean variables to factor variables.

| **Boolean *Microhabitat* variables** | | | | **Factor *Microhabitat* variable** | **N** |
| --- | --- | --- | --- | --- | --- |
| ***Microhab_Fos*** | ***Microhab_Ter*** | ***Microhab_Aqu*** | ***Microhab_Arb*** | **New level** |  |
| 0 | 0 | 0 | 1 | *Others* | 333 |
| 0 | 0 | 1 | 0 | *Others* | 100 |
| 0 | 0 | 1 | 1 | *Generalist* | 91 |
| 0 | 1 | 0 | 0 | *Generalist* | 1110 |
| 0 | 1 | 0 | 1 | *Others* | 919 |
| 0 | 1 | 1 | 0 | *Semiaquatic* | 2268 |
| 0 | 1 | 1 | 1 | *Generalist* | 1244 |
| 1 | 0 | 0 | 0 | *Others* | 39 |
| 1 | 0 | 0 | 1 | *Generalist* | 2 |
| 1 | 0 | 1 | 0 | *Generalist* | 51 |
| 1 | 1 | 0 | 0 | *Others* | 75 |
| 1 | 1 | 0 | 1 | *Generalist* | 8 |
| 1 | 1 | 1 | 0 | *Generalist* | 513 |
| 1 | 1 | 1 | 1 | *Generalist* | 3 |

**Table A9.** Variable selection for each model. We report if a variable was included in the final model (Included) or the reason/process for exclusion: excluded by collinearity when calculating variance inflation factor (VIF), stepwise backward model selection procedure by AIC (BS), variable included in their factor version (FV), variable included in their binary version (BV), phylogenetic variables excluded in PGLS models (EP), variable excluded in a model selection approach according to their importance (SI), variable excluded due to unbalanced number of species causing problems to fit and validate models (UN). Notice that in the variable list are included variables in the factor and in their binary version which were used for algorithms using numeric and factorial databases respectively (Indicated just below the name of each model algorithm), such as the factor variable *Microhab_rec* and their binary versions *Microhabitat_Aqu, Microhabitat_Arb, Microhabitat_Fos and Microhabitat_Ter*, those variables are mutually exclusive.

|  | **Anura** | | | | **Caudata** | | | | **Gymnophiona** | | | |
| --- | --- | --- | --- | --- | --- | --- | --- | --- | --- | --- | --- | --- |
| **Variable Group** | **CLM** | **PGLS** | **RF** | **NN** | **CLM** | **PGLS** | **RF** | **NN** | **CLM** | **PGLS** | **RF** | **NN** |
| *Variable* | *Numeric* | *Factorial* | *Factorial* | *Numeric* | *Numeric* | *Factorial* | *Factorial* | *Numeric* | *Numeric* | *Factorial* | *Factorial* | *Numeric* |
| **Land cover/Land cover change** |  |  |  |  |  |  |  |  |  |  |  |  |
| *Urbanization* | Included | Included | Included | Included | BS | BS | Included | Included | BS | BS | Included | Included |
| *Agriculture* | Included | Included | SI | Included | BS | BS | SI | Included | Included | Included | SI | Included |
| *Urbanization_Change* | VIF | VIF | VIF | VIF | VIF | VIF | VIF | VIF | VIF | VIF | VIF | VIF |
| *Agriculture_Change* | BS | BS | SI | Included | BS | BS | SI | Included | BS | BS | SI | Included |
| *Human_density* | Included | Included | Included | Included | BS | Included | SI | Included | Included | BS | SI | Included |
| *Accessibility* | Included | Included | Included | Included | BS | BS | SI | Included | Included | BS | Included | Included |
| **Spatial context** |  |  |  |  |  |  |  |  |  |  |  |  |
| *Realm* | BV | Included | Included | BV | BV | Included | Included | BV | BV | BS | SI | BV |
| *Realm_1* | Included | FV | FV | Included | UN | FV | FV | UN | UN | FV | FV | UN |
| *Realm_3* | Included | FV | FV | Included | UN | FV | FV | UN | VIF | FV | FV | VIF |
| *Realm_4* | BS | FV | FV | Included | BS | FV | FV | Included | UN | FV | FV | Included |
| *Realm_5* | Included | FV | FV | Included | Included | FV | FV | Included | UN | FV | FV | UN |
| *Realm_6* | VIF | FV | FV | VIF | Included | FV | FV | Included | Included | FV | FV | Included |
| **Climate** |  |  |  |  |  |  |  |  |  |  |  |  |
| *Climate_1* | Included | BS | SI | Included | Included | Included | Included | Included | Included | Included | SI | Included |
| *Climate_2* | Included | Included | Included | Included | Included | Included | Included | Included | Included | BS | Included | Included |
| *Climate_3* | Included | Included | Included | Included | BS | BS | SI | Included | Included | BS | SI | Included |
| *Climate_4* | Included | Included | Included | Included | Included | Included | Included | Included | Included | Included | SI | Included |
| **Climate Change** |  |  |  |  |  |  |  |  |  |  |  |  |
| *Climate_Change_1* | Included | BS | SI | Included | BS | Included | SI | Included | VIF | VIF | VIF | VIF |
| *Climate_Change_2* | Included | Included | Included | Included | Included | BS | SI | Included | BS | Included | SI | Included |
| *Climate_Change_3* | BS | Included | SI | Included | BS | Included | SI | Included | Included | Included | SI | Included |
| *Climate_Change_4* | BS | Included | SI | Included | Included | Included | SI | Included | Included | BS | SI | Included |
| *Climate_Change_5* | Included | Included | Included | Included | Included | BS | SI | Included | BS | Included | Included | Included |
| **Ecological traits** |  |  |  |  |  |  |  |  |  |  |  |  |
| *Body_size* | Included | Included | SI | Included | Included | BS | SI | Included | Included | BS | SI | Included |
| *Brood_size* | Included | BS | SI | Included | Included | Included | SI | Included | Included | BS | SI | Included |
| *Parity* | BV | BS | SI | BV | BV | BS | SI | BV | BV | Included | SI | BV |
| *Parity_Direct* | VIF | FV | FV | VIF | VIF | FV | FV | VIF | VIF | FV | FV | VIF |
| *Parity_Larvae* | BS | FV | FV | Included | Included | FV | FV | Included | VIF | FV | FV | VIF |
| *Parity_Viviparous* | BS | FV | FV | Included | BS | FV | FV | Included | UN | FV | FV | Included |
| *Generalist* | BV | Included | SI | BV | BV | Included | SI | BV | BV | BS | SI | BV |
| *Specialist_NoForest* | BS | FV | FV | Included | Included | FV | FV | Included | UN | FV | FV | UN |
| *Specialist_Forest* | Included | FV | FV | Included | BS | FV | FV | Included | VIF | FV | FV | VIF |
| *Generalist_NoForest* | VIF | FV | FV | VIF | VIF | FV | FV | VIF | VIF | FV | FV | VIF |
| *Generalist_Forest* | Included | FV | FV | Included | Included | FV | FV | Included | UN | FV | FV | Included |
| *Microhab_rec* | BV | Included | SI | BV | BV | BS | SI | BV | BV | BS | SI | BV |
| *Microhabitat_Aqu* | Included | FV | FV | Included | Included | FV | FV | Included | UN | FV | FV | Included |
| *Microhabitat_Arb* | BS | FV | FV | Included | BS | FV | FV | Included | UN | FV | FV | UN |
| *Microhabitat_Fos* | BS | FV | FV | Included | BS | FV | FV | Included | UN | FV | FV | UN |
| *Microhabitat_Ter* | Included | FV | FV | Included | BS | FV | FV | Included | UN | FV | FV | Included |
| **Range/Spatial configuration** |  |  |  |  |  |  |  |  |  |  |  |  |
| *Range_area* | Included | Included | Included | Included | Included | Included | Included | Included | Included | Included | Included | Included |
| *Range_fragments* | Included | BS | SI | Included | BS | BS | SI | Included | Included | BS | SI | Included |
| *Range_Circularity* | Included | Included | Included | Included | BS | Included | Included | Included | BS | BS | SI | Included |
| *Range_heterogeinity* | Included | Included | SI | Included | Included | BS | SI | Included | Included | Included | SI | Included |
| **Phylogeny** |  |  |  |  |  |  |  |  |  |  |  |  |
| *Phylogeny_1* | Included | EP | SI | Included | Included | EP | SI | Included | BS | EP | SI | Included |
| *Phylogeny_2* | VIF | EP | VIF | VIF | VIF | EP | VIF | VIF | VIF | EP | VIF | VIF |
| *Phylogeny_3* | Included | EP | SI | Included | VIF | EP | VIF | VIF | VIF | EP | VIF | VIF |
| *Phylogeny_4* | Included | EP | SI | Included | Included | EP | SI | Included | VIF | EP | VIF | VIF |
| N variables in the final model | 27 | 18 | 11 | 36 | 18 | 12 | 7 | 34 | 15 | 9 | 5 | 26 |

**Table A10.** Variable importance for each model, for the averaged importance within each order and for the averaged importance for the three orders and four model algorithms (Amphibians). Variables are ordered using the averaged importance for the three orders and four model algorithms. The ten most important variables within Anura, Caudata, Gymnophiona and Amphibians are shaded in grey. To calculate the averaged variable importance by order and for Amphibians, we considered NA values (which represent variables not included in the model) as 0.

|  | **Anura** | | | | **Caudata** | | | | **Gymnophiona** | | | | **Anura** | **Caudata** | **Gymnophiona** | **Amphibians** |
| --- | --- | --- | --- | --- | --- | --- | --- | --- | --- | --- | --- | --- | --- | --- | --- | --- |
| **Variable** | **CLM** | **PGLS** | **RF** | **NN** | **CLM** | **PGLS** | **RF** | **NN** | **CLM** | **PGLS** | **RF** | **NN** |  |  |  |  |
| ***Range_area*** | 0.227 | 0.238 | 0.322 | 0.054 | 0.187 | 0.305 | 0.373 | 0.033 | 0.061 | 0.222 | 0.468 | 0.043 | 0.210 | 0.224 | 0.198 | 0.211 |
| ***Realm*** | 0.065 | 0.123 | 0.068 | 0.110 | 0.131 | 0.095 | 0.116 | 0.086 | 0.039 | NA | NA | 0.072 | 0.091 | 0.107 | 0.028 | 0.075 |
| ***Climate_2*** | 0.034 | 0.079 | 0.092 | 0.028 | 0.071 | 0.045 | 0.155 | 0.036 | 0.059 | NA | 0.110 | 0.038 | 0.058 | 0.077 | 0.052 | 0.062 |
| ***Habitat_specialist*** | 0.076 | 0.092 | NA | 0.075 | 0.119 | 0.076 | NA | 0.089 | NA | NA | NA | 0.040 | 0.061 | 0.071 | 0.010 | 0.047 |
| ***Urbanization*** | 0.013 | 0.032 | 0.106 | 0.028 | NA | NA | 0.081 | 0.025 | NA | NA | 0.215 | 0.037 | 0.045 | 0.027 | 0.063 | 0.045 |
| ***Climate_4*** | 0.052 | 0.063 | 0.045 | 0.028 | 0.041 | 0.045 | 0.068 | 0.027 | 0.046 | 0.074 | NA | 0.040 | 0.047 | 0.045 | 0.040 | 0.044 |
| ***Accessibility*** | 0.103 | 0.041 | 0.081 | 0.038 | NA | NA | NA | 0.031 | 0.061 | NA | 0.119 | 0.038 | 0.066 | 0.008 | 0.055 | 0.043 |
| ***Climate_1*** | 0.013 | NA | NA | 0.028 | 0.062 | 0.062 | 0.101 | 0.030 | 0.082 | 0.074 | NA | 0.039 | 0.010 | 0.064 | 0.049 | 0.041 |
| ***Climate_Change_5*** | 0.026 | 0.043 | 0.042 | 0.031 | 0.031 | NA | NA | 0.031 | NA | 0.083 | 0.088 | 0.040 | 0.036 | 0.016 | 0.053 | 0.035 |
| ***Microhabitat*** | 0.046 | 0.020 | NA | 0.100 | 0.054 | NA | NA | 0.115 | NA | NA | NA | 0.074 | 0.041 | 0.042 | 0.018 | 0.034 |
| ***Climate_Change_2*** | 0.014 | 0.036 | 0.061 | 0.031 | 0.035 | NA | NA | 0.032 | NA | 0.153 | NA | 0.040 | 0.036 | 0.017 | 0.048 | 0.033 |
| ***Range_Circularity*** | 0.015 | 0.037 | 0.055 | 0.030 | NA | 0.076 | 0.107 | 0.032 | NA | NA | NA | 0.040 | 0.034 | 0.054 | 0.010 | 0.033 |
| ***Human_density*** | 0.014 | 0.061 | 0.058 | 0.029 | NA | 0.111 | NA | 0.020 | 0.036 | NA | NA | 0.041 | 0.041 | 0.033 | 0.019 | 0.031 |
| ***Range_heterogeinity*** | 0.027 | 0.014 | NA | 0.026 | 0.039 | NA | NA | 0.037 | 0.070 | 0.090 | NA | 0.040 | 0.017 | 0.019 | 0.050 | 0.028 |
| ***Parity*** | NA | NA | NA | 0.036 | 0.038 | NA | NA | 0.060 | NA | 0.154 | NA | 0.040 | 0.009 | 0.024 | 0.049 | 0.027 |
| ***Climate_Change_3*** | NA | 0.023 | NA | 0.028 | NA | 0.047 | NA | 0.027 | 0.081 | 0.073 | NA | 0.036 | 0.013 | 0.018 | 0.048 | 0.026 |
| ***Brood_size*** | 0.032 | NA | NA | 0.025 | 0.023 | 0.054 | NA | 0.030 | 0.110 | NA | NA | 0.037 | 0.014 | 0.027 | 0.037 | 0.026 |
| ***Climate_3*** | 0.021 | 0.038 | 0.071 | 0.032 | NA | NA | NA | 0.030 | 0.070 | NA | NA | 0.037 | 0.041 | 0.008 | 0.027 | 0.025 |
| ***Body_size*** | 0.041 | 0.018 | NA | 0.028 | 0.061 | NA | NA | 0.032 | 0.078 | NA | NA | 0.039 | 0.022 | 0.023 | 0.029 | 0.025 |
| ***Climate_Change_4*** | NA | 0.025 | NA | 0.029 | 0.049 | 0.039 | NA | 0.034 | 0.082 | NA | NA | 0.036 | 0.014 | 0.031 | 0.029 | 0.025 |
| ***Agriculture*** | 0.021 | 0.018 | NA | 0.033 | NA | NA | NA | 0.027 | 0.039 | 0.076 | NA | 0.037 | 0.018 | 0.007 | 0.038 | 0.021 |
| ***Range_fragments*** | 0.022 | NA | NA | 0.017 | NA | NA | NA | 0.021 | 0.087 | NA | NA | 0.037 | 0.010 | 0.005 | 0.031 | 0.015 |
| ***Phylogeny_1*** | 0.056 | NA | NA | 0.027 | 0.030 | NA | NA | 0.030 | NA | NA | NA | 0.038 | 0.021 | 0.015 | 0.009 | 0.015 |
| ***Climate_Change_1*** | 0.031 | NA | NA | 0.026 | NA | 0.044 | NA | 0.031 | NA | NA | NA | NA | 0.014 | 0.019 | NA | 0.011 |
| ***Phylogeny_4*** | 0.038 | NA | NA | 0.025 | 0.030 | NA | NA | 0.028 | NA | NA | NA | NA | 0.016 | 0.014 | NA | 0.010 |
| ***Agriculture_Change*** | NA | NA | NA | 0.032 | NA | NA | NA | 0.027 | NA | NA | NA | 0.041 | 0.008 | 0.007 | 0.010 | 0.008 |
| ***Phylogeny_3*** | 0.012 | NA | NA | 0.025 | NA | NA | NA | NA | NA | NA | NA | NA | 0.009 | NA | NA | 0.003 |
| ***Urbanization_Change*** | NA | NA | NA | NA | NA | NA | NA | NA | NA | NA | NA | NA | NA | NA | NA | NA |
| ***Phylogeny_2*** | NA | NA | NA | NA | NA | NA | NA | NA | NA | NA | NA | NA | NA | NA | NA | NA |

**Table A11.** Coefficients, standard error (SE), T-values (coefficient/SE) and P-value for each of the variables included in the cumulative link model (CLM) predicting extinction risk in Anura.

| **Variable** | **Coefficient** | **Std. Error** | **T-values (coefficient/SE)** | **P-value** |
| --- | --- | --- | --- | --- |
| ***Phylogeny_1*** | 0.167 | 0.022 | 7.76 | 0 |
| ***Phylogeny_3*** | -0.043 | 0.025 | -1.688 | 0.091 |
| ***Phylogeny_4*** | -0.155 | 0.029 | -5.289 | 0 |
| ***Urbanization*** | -0.04 | 0.023 | -1.754 | 0.079 |
| ***Agriculture*** | 0.055 | 0.019 | 2.882 | 0.004 |
| ***Human_density*** | 0.044 | 0.022 | 1.991 | 0.046 |
| ***Accessibility*** | -0.215 | 0.015 | -14.207 | 0 |
| ***Body_size*** | 0.133 | 0.023 | 5.691 | 0 |
| ***Brood_size*** | -0.121 | 0.028 | -4.36 | 0 |
| ***Range_Circularity*** | 0.041 | 0.02 | 2.111 | 0.035 |
| ***Range_fragments*** | -0.938 | 0.316 | -2.973 | 0.003 |
| ***Range_heterogeinity*** | 0.065 | 0.018 | 3.66 | 0 |
| ***Climate_1*** | 0.043 | 0.024 | 1.771 | 0.077 |
| ***Climate_2*** | 0.117 | 0.025 | 4.615 | 0 |
| ***Climate_3*** | -0.07 | 0.024 | -2.923 | 0.003 |
| ***Climate_4*** | 0.126 | 0.018 | 7.173 | 0 |
| ***Climate_Change_1*** | 0.113 | 0.026 | 4.301 | 0 |
| ***Climate_Change_2*** | -0.059 | 0.029 | -1.994 | 0.046 |
| ***Climate_Change_5*** | 0.055 | 0.015 | 3.642 | 0 |
| ***Range_area*** | -1.031 | 0.033 | -31.294 | 0 |
| ***Microhab_Ter*** | -0.101 | 0.058 | -1.739 | 0.082 |
| ***Microhab_Aqu*** | 0.162 | 0.036 | 4.55 | 0 |
| ***Generalist_Forest*** | 0.306 | 0.055 | 5.579 | 0 |
| ***Specialist_Forest*** | 0.319 | 0.066 | 4.861 | 0 |
| ***Realm_Australasian*** | -0.259 | 0.072 | -3.588 | 0 |
| ***Realm_Afrotropical*** | 0.187 | 0.057 | 3.309 | 0.001 |
| ***Realm_Neartic*** | 0.292 | 0.143 | 2.039 | 0.041 |

**Table A12.** Coefficients, standard error (SE), t-values (coefficient/SE) and p-value for each of the variables included in the phylogenetic generalized least squares model (PGLS) predicting extinction risk in Anura.

| **Variable** | **Coefficient** | **Std. Error** | **T-values (coefficient/SE)** | **P-value** |
| --- | --- | --- | --- | --- |
| ***Urbanization*** | 0.085 | 0.024 | 3.563 | 0 |
| ***Agrigulture*** | -0.036 | 0.018 | -2.017 | 0.044 |
| ***Human_density*** | -0.151 | 0.022 | -6.813 | 0 |
| ***Accessibility*** | -0.098 | 0.021 | -4.578 | 0 |
| ***Body_size*** | 0.047 | 0.024 | 1.97 | 0.049 |
| ***Range_circularity*** | 0.074 | 0.018 | 4.109 | 0 |
| ***Range_heterogeinity*** | -0.024 | 0.015 | -1.543 | 0.123 |
| ***Climate_2*** | 0.193 | 0.022 | 8.769 | 0 |
| ***Climate_3*** | -0.102 | 0.024 | -4.284 | 0 |
| ***Climate_4*** | 0.139 | 0.02 | 7.011 | 0 |
| ***Climate_Change_2*** | 0.113 | 0.028 | 4.04 | 0 |
| ***Climate_Change_3*** | -0.067 | 0.026 | -2.553 | 0.011 |
| ***Climate_Change_4*** | 0.06 | 0.021 | 2.832 | 0.005 |
| ***Climate_Change_5*** | 0.086 | 0.018 | 4.789 | 0 |
| ***Range_area*** | -0.668 | 0.025 | -26.544 | 0 |
| ***Realm_Afrotropical*** | 0.479 | 0.284 | 1.685 | 0.092 |
| ***Realm_Indomalayan*** | 0.634 | 0.177 | 3.581 | 0 |
| ***Realm_Neartic*** | 1.017 | 0.35 | 2.906 | 0.004 |
| ***Realm_Neotropical*** | 0.678 | 0.315 | 2.156 | 0.031 |
| ***Realm_Paleartic*** | 0.694 | 0.207 | 3.362 | 0.001 |
| ***Generalist_Forest*** | 0.125 | 0.046 | 2.71 | 0.007 |
| ***Specialist_NonForest*** | 0.268 | 0.099 | 2.71 | 0.007 |
| ***Specialist_Forest*** | 0.317 | 0.065 | 4.886 | 0 |
| ***Microhabitat_Others*** | 0.093 | 0.044 | 2.099 | 0.036 |
| ***Microhabitat_Semiaquatic*** | -0.005 | 0.039 | -0.124 | 0.901 |

**Figure A2.** Partial dependence functions (red line) for each of the variables included in the random forest model explaining extinction risk for Anurans. In grey we show the individual conditional expectation (ICE) curves for each data used to fit the model.

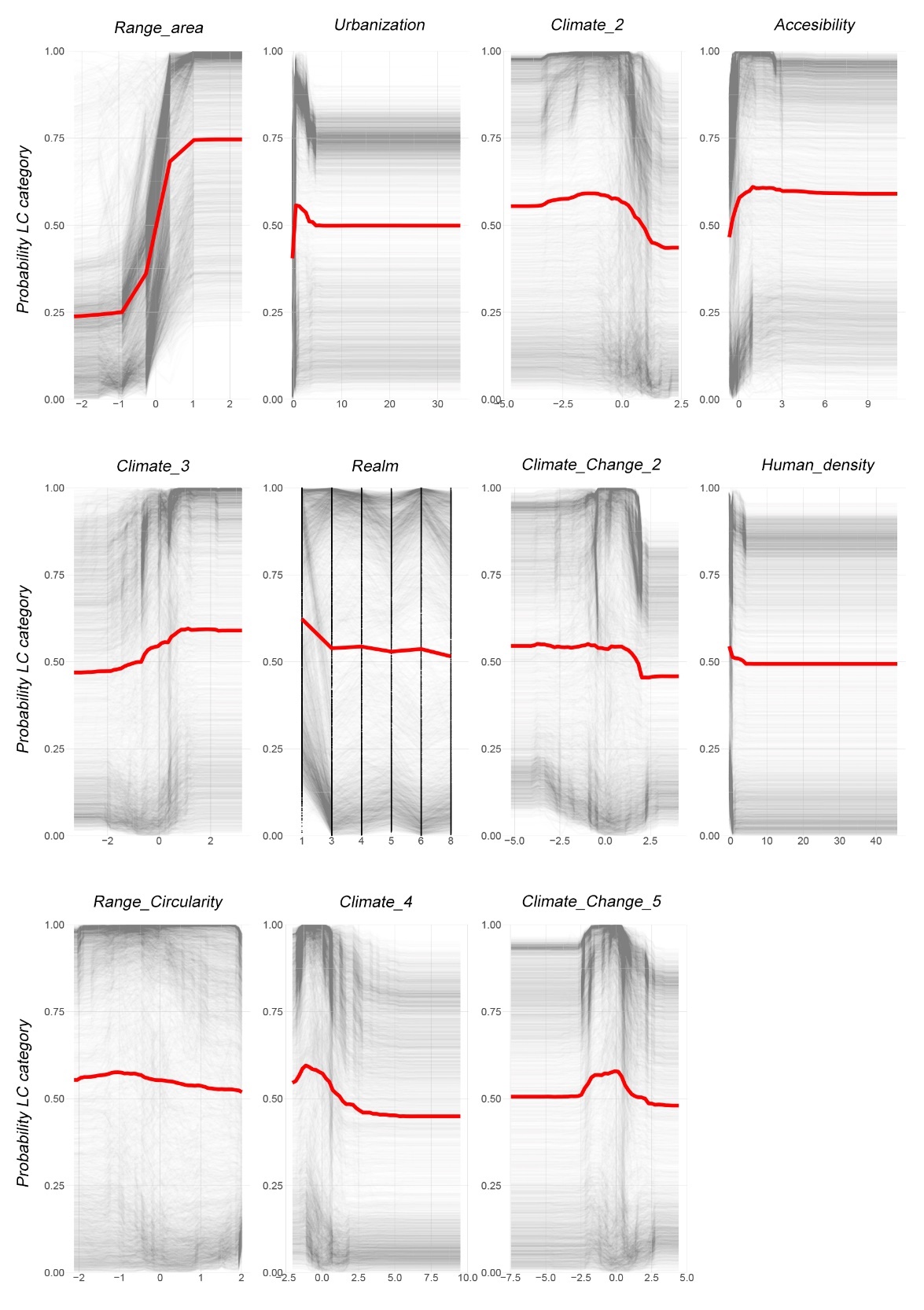

**Figure A3.** Partial dependence plots for each of the variables included in the neural network model (NN) explaining extinction risk for Anura. Red line indicates the marginal effect of each variable on the extinction risk (measured as a continuous variable ranging from 1 for Least Concern to 5 for Critically Endangered).

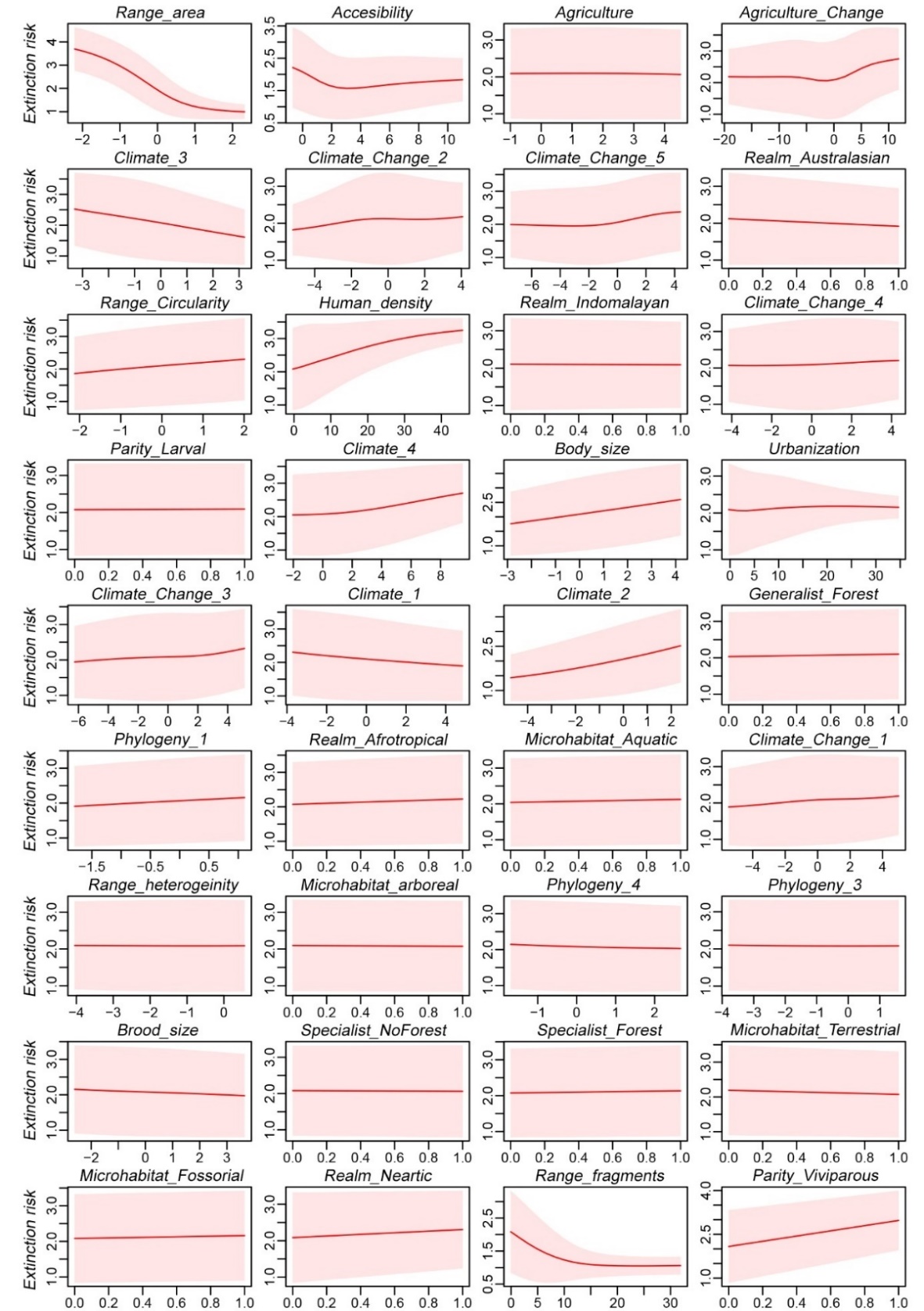

**Table A13.** Coefficients, standard error (SE), T-values (coefficient/SE) and P-value for each of the variables included in the cumulative link model (CLM) predicting extinction risk in Caudata.

| **Variable** | **Coefficient** | **Std. Error** | **T-values (coefficient/SE)** | **P-value** |
| --- | --- | --- | --- | --- |
| ***Phylogeny_1*** | -0.172 | 0.092 | -1.881 | 0.06 |
| ***Phylogeny_4*** | -0.215 | 0.112 | -1.921 | 0.055 |
| ***Body_size*** | 0.238 | 0.062 | 3.842 | 0 |
| ***Brood_size*** | -0.091 | 0.063 | -1.444 | 0.149 |
| ***Range_heterogeinity*** | 0.124 | 0.051 | 2.452 | 0.014 |
| ***Climate_1*** | 0.352 | 0.089 | 3.943 | 0 |
| ***Climate_2*** | 0.367 | 0.081 | 4.524 | 0 |
| ***Climate_4*** | 0.153 | 0.06 | 2.577 | 0.01 |
| ***Climate_Change_2*** | -0.132 | 0.061 | -2.187 | 0.029 |
| ***Climate_Change_4*** | -0.181 | 0.058 | -3.111 | 0.002 |
| ***Climate_Change_5*** | -0.112 | 0.056 | -1.994 | 0.046 |
| ***Range_area*** | -1.179 | 0.1 | -11.813 | 0 |
| ***Microhabitat_Aqu*** | 0.537 | 0.158 | 3.394 | 0.001 |
| ***Generalist_Forest*** | -0.623 | 0.123 | -5.045 | 0 |
| ***Specialist_NonForest*** | -0.579 | 0.231 | -2.502 | 0.012 |
| ***Parity_Larve*** | 0.446 | 0.187 | 2.393 | 0.017 |
| ***Realm_Neartic*** | -0.939 | 0.184 | -5.091 | 0 |
| ***Realm_Neotropical*** | 0.782 | 0.245 | 3.193 | 0.001 |

**Table A14.** Coefficients, standard error (SE), t-values (coefficient/SE) and p-value for each of the variables included in the phylogenetic generalized least squares model (PGLS) predicting extinction risk in Caudata.

| **Variable** | **Coefficient** | **Std. Error** | **T-values (coefficient/SE)** | **P-value** |
| --- | --- | --- | --- | --- |
| ***Human_density*** | 0.129 | 0.033 | 3.934 | 0 |
| ***Brood_size*** | -0.118 | 0.061 | -1.927 | 0.055 |
| ***Range_circularity*** | 0.123 | 0.046 | 2.687 | 0.007 |
| ***Climate_1*** | 0.171 | 0.077 | 2.208 | 0.028 |
| ***Climate_2*** | 0.097 | 0.06 | 1.617 | 0.107 |
| ***Climate_4*** | 0.101 | 0.063 | 1.602 | 0.11 |
| ***Climate_Change_1*** | 0.094 | 0.059 | 1.576 | 0.116 |
| ***Climate_Change_3*** | -0.122 | 0.073 | -1.663 | 0.097 |
| ***Climate_Change_4*** | -0.095 | 0.068 | -1.399 | 0.162 |
| ***Range_area*** | -0.714 | 0.066 | -10.857 | 0 |
| ***Realm_Neartic*** | -0.701 | 0.825 | -0.849 | 0.396 |
| ***Realm_Neotropical*** | 0.212 | 0.876 | 0.242 | 0.809 |
| ***Realm_Paleartic*** | -0.617 | 0.269 | -2.296 | 0.022 |
| ***Generalist_Forest*** | -0.27 | 0.219 | -1.23 | 0.219 |
| ***Specialist_NonForest*** | -0.314 | 0.239 | -1.315 | 0.189 |
| ***Specialist_Forest*** | -0.036 | 0.239 | -0.152 | 0.879 |

**Figure A4.** Partial dependence functions (red line) for each of the variables included in the random forest model explaining extinction risk for Caudata. In grey we show the individual conditional expectation (ICE) curves for each data used to fit the model.

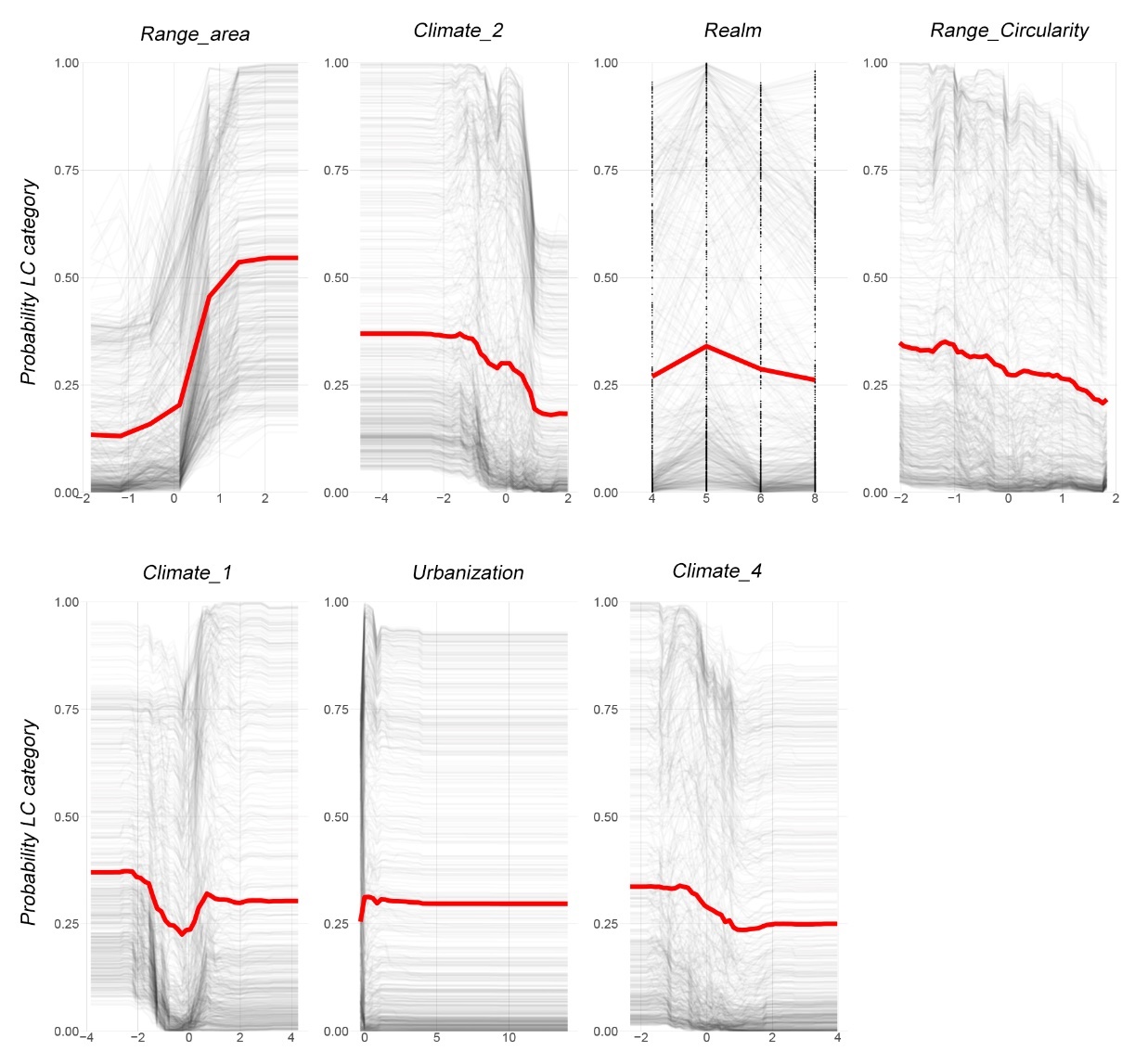

**Figure A5.** Partial dependence plots for each of the variables included in the neural network model (NN) explaining extinction risk for Caudata. Red line indicates the marginal effect of each variable on the extinction risk (measured as a continuous variable ranging from 1 for Least Concern to 5 for Critically Endangered).

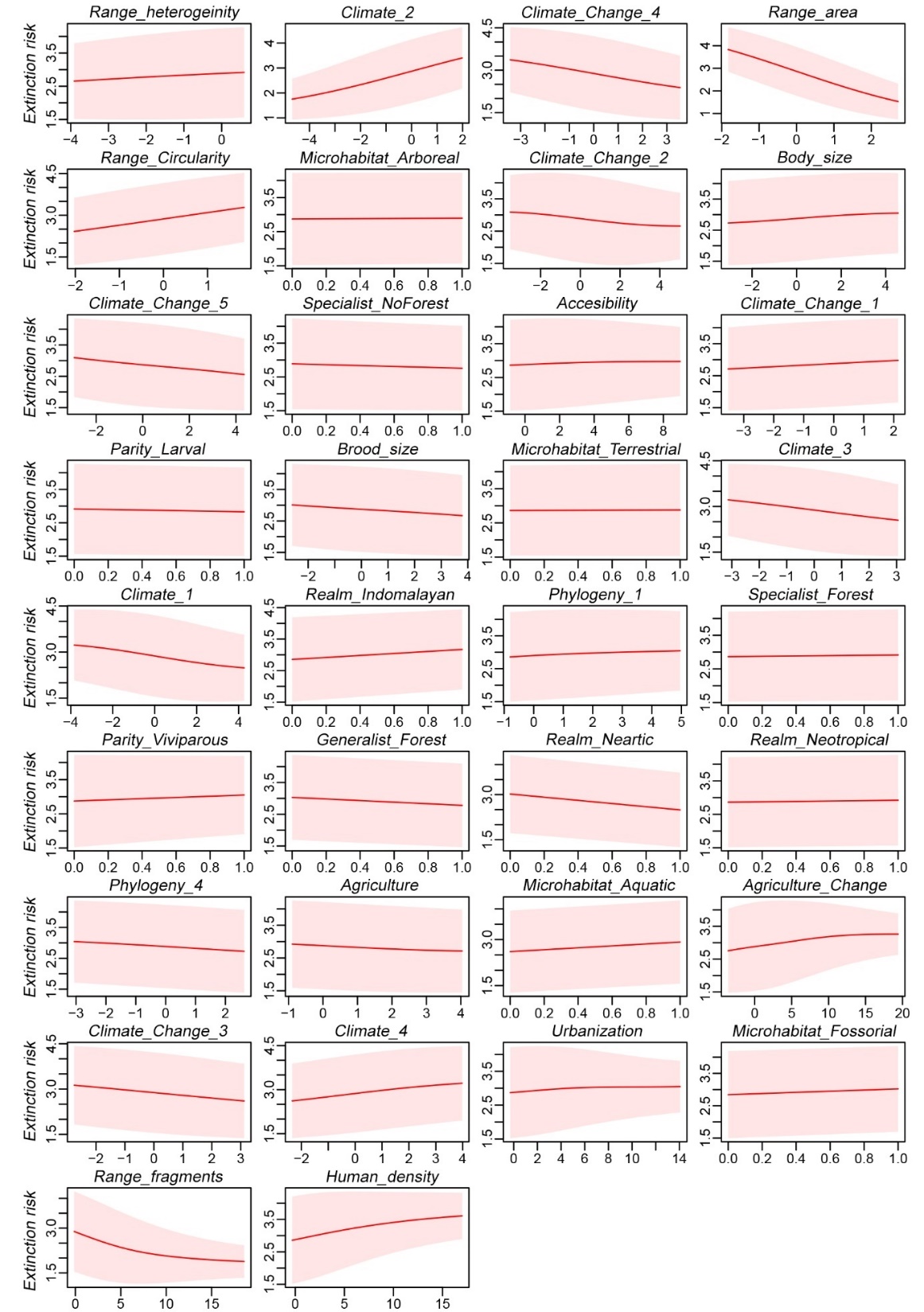

**Table A15.** Coefficients, standard error (SE), T-values (coefficient/SE) and P-value for each of the variables included in the cumulative link model (CLM) predicting extinction risk in Gymnophiona.

| **Variable** | **Coefficient** | **Std. Error** | **T-values (coefficient/SE)** | **P-value** |
| --- | --- | --- | --- | --- |
| ***Agriculture*** | 0.434 | 0.28 | 1.553 | 0.12 |
| ***Human_density*** | -0.341 | 0.234 | -1.457 | 0.145 |
| ***Accesibility*** | -1.012 | 0.415 | -2.437 | 0.015 |
| ***Body_size*** | -0.791 | 0.252 | -3.14 | 0.002 |
| ***Brood_size*** | 1.392 | 0.316 | 4.405 | 0 |
| ***Range_fragments*** | -30.402 | 8.709 | -3.491 | 0 |
| ***Range_heterogeinity*** | -1.302 | 0.461 | -2.827 | 0.005 |
| ***Climate_1*** | 1.16 | 0.351 | 3.306 | 0.001 |
| ***Climate_2*** | 1.098 | 0.464 | 2.364 | 0.018 |
| ***Climate_3*** | -1.188 | 0.424 | -2.804 | 0.005 |
| ***Climate_4*** | 0.559 | 0.302 | 1.849 | 0.064 |
| ***Climate_Change_3*** | 1.046 | 0.32 | 3.271 | 0.001 |
| ***Climate_Change_4*** | -1.036 | 0.316 | -3.279 | 0.001 |
| ***Range_area*** | -1.19 | 0.489 | -2.431 | 0.015 |
| ***Realm_Neotropical*** | 1.13 | 0.725 | 1.559 | 0.119 |

**Table A16.** Coefficients, standard error (SE), t-values (coefficient/SE) and p-value for each of the variables included in the phylogenetic generalized least squares model (PGLS) predicting extinction risk in Gymnophiona.

| **Variable** | **Coefficient** | **Std. Error** | **T-values (coefficient/SE)** | **P-value** |
| --- | --- | --- | --- | --- |
| ***Agriculture*** | 0.208 | 0.129 | 1.603 | 0.113 |
| ***Range_heterogeinity*** | 0.142 | 0.075 | 1.899 | 0.062 |
| ***Climate_1*** | -0.148 | 0.095 | -1.567 | 0.121 |
| ***Climate_4*** | -0.164 | 0.104 | -1.567 | 0.121 |
| ***Climate_Change_2*** | 0.429 | 0.133 | 3.218 | 0.002 |
| ***Climate_Change_3*** | -0.203 | 0.131 | -1.543 | 0.127 |
| ***Climate_Change_5*** | -0.205 | 0.117 | -1.75 | 0.084 |
| ***Range_area*** | -0.541 | 0.115 | -4.691 | 0 |
| ***Parity_Larve*** | -0.367 | 0.385 | -0.952 | 0.344 |
| ***Parity_Vivipary*** | -0.908 | 0.394 | -2.305 | 0.024 |

**Figure A6.** Partial dependence functions (red line) for each of the variables included in the random forest model explaining extinction risk for Gymnophiona. In grey we show the individual conditional expectation (ICE) curves for each data used to fit the model.

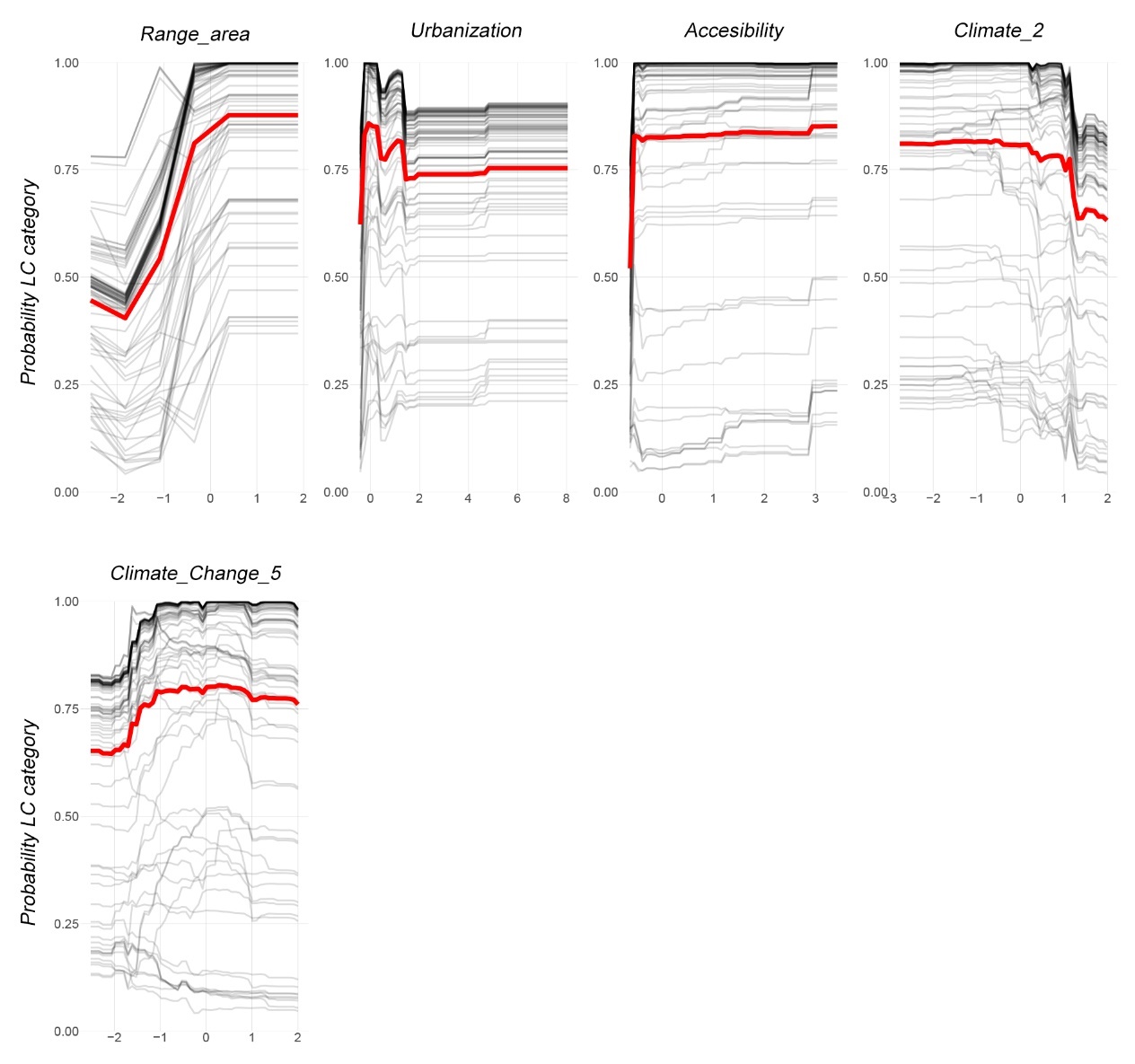

**Figure A7.** Partial dependence plots for each of the variables included in the neural network model (NN) explaining extinction risk for Gymnophiona. Red line indicates the marginal effect of each variable on the extinction risk (measured as a continuous variable ranging from 1 for Least Concern to 5 for Critically Endangered).

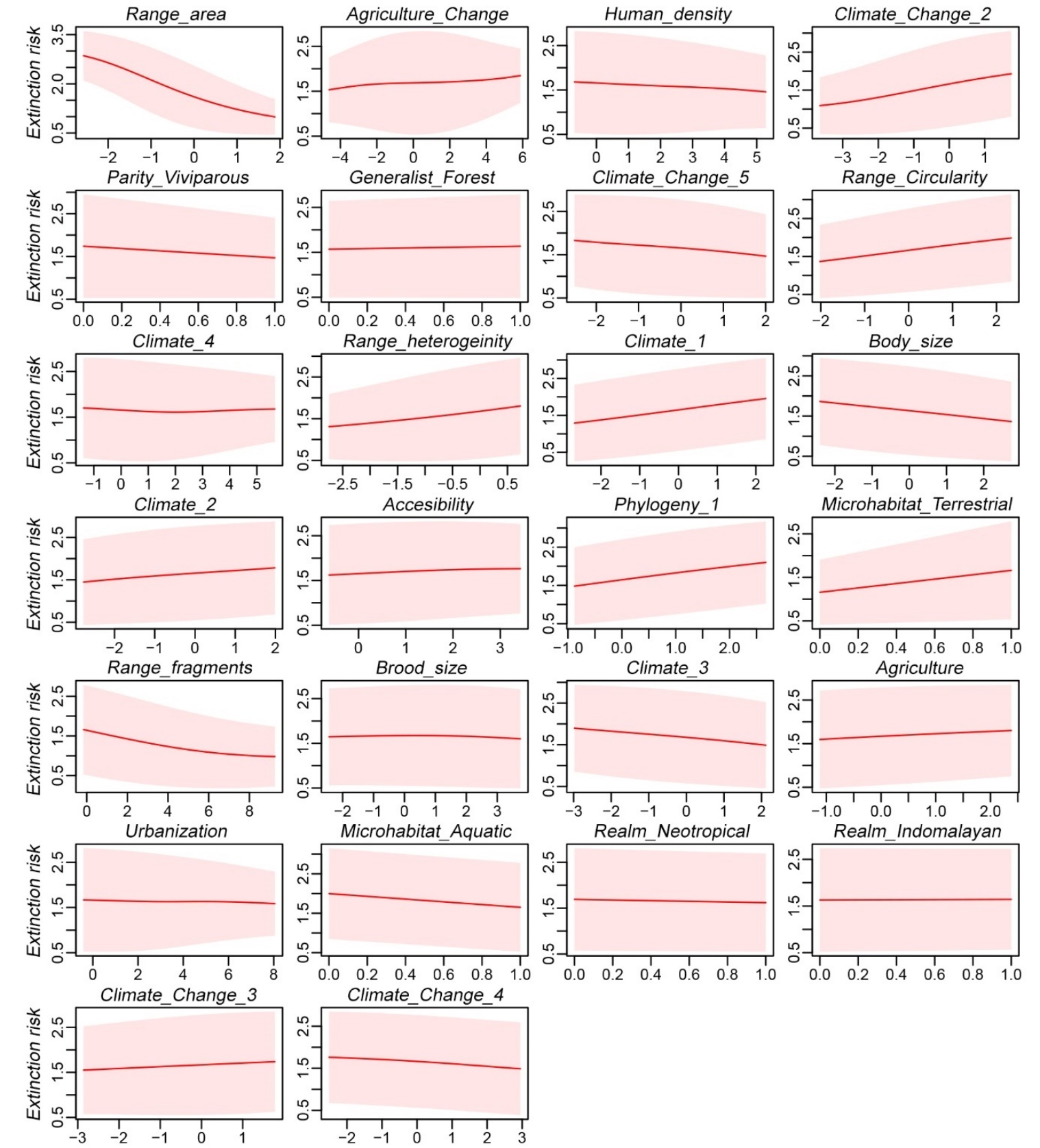

**Table A17.** Result for family block validation using as response variable Red List categories. We show the accuracy (rate of correct classification) the true skill statistic (TSS = sensitivity + specificity -1), the sensitivity, the specificity, and the mean absolute error (Error) for each category within each of the 12 fitted models (3 orders x 4 model algorithms).

| **Order** | **Algorithm** | **Category** | **Accuracy** | **TSS** | **Sensitivity** | **Specificity** | **Error** |
| --- | --- | --- | --- | --- | --- | --- | --- |
| ANURA | CLM | LC | 0.78 | 0.59 | 0.67 | 0.92 | 0.56 |
| ANURA | CLM | NT | 0.77 | 0.19 | 0.38 | 0.80 | 0.95 |
| ANURA | CLM | VU | 0.84 | 0.19 | 0.28 | 0.91 | 0.96 |
| ANURA | CLM | EN | 0.83 | 0.32 | 0.40 | 0.92 | 0.82 |
| ANURA | CLM | CR | 0.90 | 0.61 | 0.69 | 0.92 | 0.52 |
| ANURA | PGLS | LC | 0.68 | 0.42 | 0.47 | 0.96 | 0.66 |
| ANURA | PGLS | NT | 0.64 | 0.20 | 0.55 | 0.64 | 0.51 |
| ANURA | PGLS | VU | 0.73 | 0.17 | 0.40 | 0.77 | 0.66 |
| ANURA | PGLS | EN | 0.78 | 0.04 | 0.12 | 0.92 | 1.16 |
| ANURA | PGLS | CR | 0.92 | 0.03 | 0.03 | 1.00 | 1.54 |
| ANURA | RF | LC | 0.84 | 0.67 | 0.91 | 0.76 | 0.23 |
| ANURA | RF | NT | 0.92 | 0.12 | 0.15 | 0.97 | 1.05 |
| ANURA | RF | VU | 0.85 | 0.25 | 0.33 | 0.92 | 1.05 |
| ANURA | RF | EN | 0.84 | 0.44 | 0.55 | 0.90 | 0.76 |
| ANURA | RF | CR | 0.93 | 0.39 | 0.41 | 0.98 | 1.04 |
| ANURA | NN | LC | 0.76 | 0.56 | 0.65 | 0.91 | 0.50 |
| ANURA | NN | NT | 0.74 | 0.12 | 0.35 | 0.77 | 0.78 |
| ANURA | NN | VU | 0.79 | 0.20 | 0.36 | 0.84 | 0.80 |
| ANURA | NN | EN | 0.81 | 0.27 | 0.37 | 0.90 | 0.86 |
| ANURA | NN | CR | 0.91 | 0.11 | 0.13 | 0.98 | 1.39 |
| CAUDATA | CLM | LC | 0.85 | 0.61 | 0.70 | 0.91 | 0.45 |
| CAUDATA | CLM | NT | 0.79 | 0.12 | 0.27 | 0.85 | 0.77 |
| CAUDATA | CLM | VU | 0.73 | 0.15 | 0.31 | 0.83 | 0.88 |
| CAUDATA | CLM | EN | 0.76 | 0.33 | 0.48 | 0.85 | 0.65 |
| CAUDATA | CLM | CR | 0.90 | 0.58 | 0.62 | 0.96 | 0.40 |
| CAUDATA | PGLS | LC | 0.76 | 0.27 | 0.29 | 0.98 | 1.05 |
| CAUDATA | PGLS | NT | 0.76 | 0.23 | 0.43 | 0.80 | 0.67 |
| CAUDATA | PGLS | VU | 0.68 | 0.22 | 0.49 | 0.73 | 0.55 |
| CAUDATA | PGLS | EN | 0.71 | 0.25 | 0.46 | 0.79 | 0.58 |
| CAUDATA | PGLS | CR | 0.89 | 0.38 | 0.40 | 0.98 | 0.66 |
| CAUDATA | RF | LC | 0.82 | 0.53 | 0.63 | 0.90 | 0.79 |
| CAUDATA | RF | NT | 0.86 | 0.13 | 0.18 | 0.94 | 1.18 |
| CAUDATA | RF | VU | 0.79 | 0.12 | 0.20 | 0.92 | 1.09 |
| CAUDATA | RF | EN | 0.78 | 0.57 | 0.80 | 0.77 | 0.23 |
| CAUDATA | RF | CR | 0.90 | 0.74 | 0.82 | 0.92 | 0.23 |
| CAUDATA | NN | LC | 0.79 | 0.37 | 0.40 | 0.96 | 0.93 |
| CAUDATA | NN | NT | 0.78 | 0.22 | 0.40 | 0.82 | 0.75 |
| CAUDATA | NN | VU | 0.73 | 0.24 | 0.45 | 0.80 | 0.62 |
| CAUDATA | NN | EN | 0.71 | 0.35 | 0.60 | 0.75 | 0.47 |
| CAUDATA | NN | CR | 0.88 | 0.32 | 0.34 | 0.99 | 0.71 |
| GYMNOPHIONA | CLM | LC | 0.69 | 0.28 | 0.73 | 0.56 | 0.46 |
| GYMNOPHIONA | CLM | NT | 0.85 | 0.20 | 0.33 | 0.87 | 0.67 |
| GYMNOPHIONA | CLM | VU | 0.89 | -0.07 | 0.00 | 0.93 | 1.50 |
| GYMNOPHIONA | CLM | EN | 0.88 | 0.42 | 0.50 | 0.92 | 1.30 |
| GYMNOPHIONA | CLM | CR | 0.99 | 0.00 | 0.00 | 1.00 | 1.00 |
| GYMNOPHIONA | PGLS | LC | 0.66 | 0.49 | 0.60 | 0.89 | 0.46 |
| GYMNOPHIONA | PGLS | NT | 0.58 | -0.40 | 0.00 | 0.60 | 1.00 |
| GYMNOPHIONA | PGLS | VU | 0.88 | 0.15 | 0.25 | 0.90 | 0.75 |
| GYMNOPHIONA | PGLS | EN | 0.88 | -0.01 | 0.00 | 0.99 | 1.70 |
| GYMNOPHIONA | PGLS | CR | 0.99 | 0.00 | 0.00 | 1.00 | 3.00 |
| GYMNOPHIONA | RF | LC | 0.85 | 0.36 | 0.97 | 0.39 | 0.03 |
| GYMNOPHIONA | RF | NT | 0.89 | 0.24 | 0.33 | 0.91 | 0.67 |
| GYMNOPHIONA | RF | VU | 0.95 | 0.00 | 0.00 | 1.00 | 2.00 |
| GYMNOPHIONA | RF | EN | 0.89 | 0.00 | 0.00 | 1.00 | 2.50 |
| GYMNOPHIONA | RF | CR | 0.99 | 0.00 | 0.00 | 1.00 | 3.00 |
| GYMNOPHIONA | NN | LC | 0.70 | 0.30 | 0.74 | 0.56 | 0.29 |
| GYMNOPHIONA | NN | NT | 0.75 | 0.42 | 0.67 | 0.75 | 0.33 |
| GYMNOPHIONA | NN | VU | 0.92 | 0.20 | 0.25 | 0.95 | 1.50 |
| GYMNOPHIONA | NN | EN | 0.89 | 0.00 | 0.00 | 1.00 | 2.10 |
| GYMNOPHIONA | NN | CR | 0.99 | 0.00 | 0.00 | 1.00 | 4.00 |

**Table A18.** Results for averaged values among categories of family block validation using as response variable Red List categories (Table A17). We show the averaged values among categories for accuracy (rate of correct classification), the true skill statistic (TSS = sensitivity + specificity -1), the sensitivity, the specificity, and the mean absolute error (Error) for each of the 12 fitted models (3 orders x 4 model algorithms).

| **Order** | **Algorithm** | **Accuracy** | **TSS** | **Error** |
| --- | --- | --- | --- | --- |
| ANURA | CLM | 0.82 | 0.38 | 0.76 |
| ANURA | PGLS | 0.8 | 0.25 | 0.87 |
| ANURA | RF | 0.75 | 0.17 | 0.91 |
| ANURA | NN | 0.88 | 0.37 | 0.83 |
| CAUDATA | CLM | 0.81 | 0.36 | 0.63 |
| CAUDATA | PGLS | 0.78 | 0.3 | 0.7 |
| CAUDATA | RF | 0.76 | 0.27 | 0.7 |
| CAUDATA | NN | 0.83 | 0.42 | 0.7 |
| GYMNOPHIONA | CLM | 0.86 | 0.17 | 0.99 |
| GYMNOPHIONA | PGLS | 0.85 | 0.18 | 1.64 |
| GYMNOPHIONA | RF | 0.8 | 0.05 | 1.38 |
| GYMNOPHIONA | NN | 0.91 | 0.12 | 1.64 |

**Table A19.** Results averaged by the number of species for family block validation using as response variable Red List categories. We show the averaged values for accuracy (rate of correct classification), the true skill statistic (TSS = sensitivity + specificity -1), the sensitivity, the specificity, and the mean absolute error (Error) for each of the 12 fitted models (3 orders x 4 model algorithms).

| **Order** | **Algorithm** | **Accuracy** | **Error** | **TSS** |
| --- | --- | --- | --- | --- |
| ANURA | CLM | 0.56 | 0.67 | 0.47 |
| ANURA | PGLS | 0.37 | 0.81 | 0.28 |
| ANURA | RF | 0.69 | 0.53 | 0.52 |
| ANURA | NN | 0.51 | 0.68 | 0.4 |
| CAUDATA | CLM | 0.52 | 0.6 | 0.4 |
| CAUDATA | PGLS | 0.4 | 0.74 | 0.27 |
| CAUDATA | RF | 0.57 | 0.66 | 0.46 |
| CAUDATA | NN | 0.45 | 0.71 | 0.32 |
| GYMNOPHIONA | CLM | 0.65 | 0.61 | 0.28 |
| GYMNOPHIONA | PGLS | 0.49 | 0.66 | 0.38 |
| GYMNOPHIONA | RF | 0.78 | 0.45 | 0.29 |
| GYMNOPHIONA | NN | 0.62 | 0.59 | 0.26 |

**Table A20.** Results for family block validation using as response variable threatened/non threatened. We show the accuracy (rate of correct classification), the true skill statistic (TSS = sensitivity + specificity -1), the sensitivity and the specificity for each of the 12 fitted models (3 orders x 4 model algorithms).

| **Order** | **Algorithm** | **Accuracy** | **TSS** | **Sensitivity** | **Specificity** |
| --- | --- | --- | --- | --- | --- |
| ANURA | CLM | 0.81 | 0.60 | 0.76 | 0.84 |
| ANURA | PGLS | 0.80 | 0.56 | 0.70 | 0.86 |
| ANURA | RF | 0.85 | 0.66 | 0.76 | 0.89 |
| ANURA | NN | 0.82 | 0.60 | 0.74 | 0.86 |
| CAUDATA | CLM | 0.84 | 0.68 | 0.84 | 0.84 |
| CAUDATA | PGLS | 0.82 | 0.59 | 0.91 | 0.68 |
| CAUDATA | RF | 0.85 | 0.66 | 0.94 | 0.72 |
| CAUDATA | NN | 0.81 | 0.60 | 0.89 | 0.70 |
| GYMNOPHIONA | CLM | 0.82 | 0.41 | 0.53 | 0.88 |
| GYMNOPHIONA | PGLS | 0.83 | 0.26 | 0.33 | 0.93 |
| GYMNOPHIONA | RF | 0.83 | 0.00 | 0.00 | 1.00 |
| GYMNOPHIONA | NN | 0.84 | 0.17 | 0.20 | 0.97 |

**Figure A8**. Performance metrics from the random block validation for four model algorithms (Cumulative Link Models [CLM], Random Forest [RF], Phylogenetic Generalised Least Square models [PGLS], Neural Network [NN]) predicting Red List categories for each of the three orders. We report the accuracy (the rate of correct classification in all categories), the mean error (absolute value of the difference between predicted and current RL categories) and the true skill statistic (TSS = Specificity + Sensitivity - 1) averaged among categories (independently of the number of species in each category) and by the number of species.

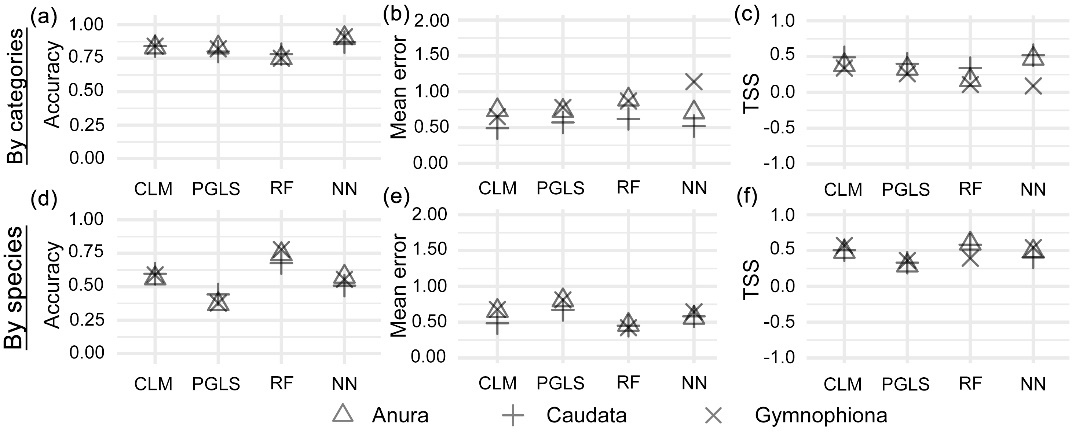

**Figure A9.** Performance metrics from the random block validation for models predicting binary (threatened/non-threatened) risk categories. Accuracy (Rate of correct classification), true skill statistic (TSS = Specificity + Sensitivity - 1), specificity (Rate of correct classification of non-threatened species), and sensitivity (Rate of correct classification of threatened species) are reported for each taxonomic order and each model algorithm. For the TSS plot, the horizontal dashed line (TSS = 0.5) indicates the value at which models are considered good.

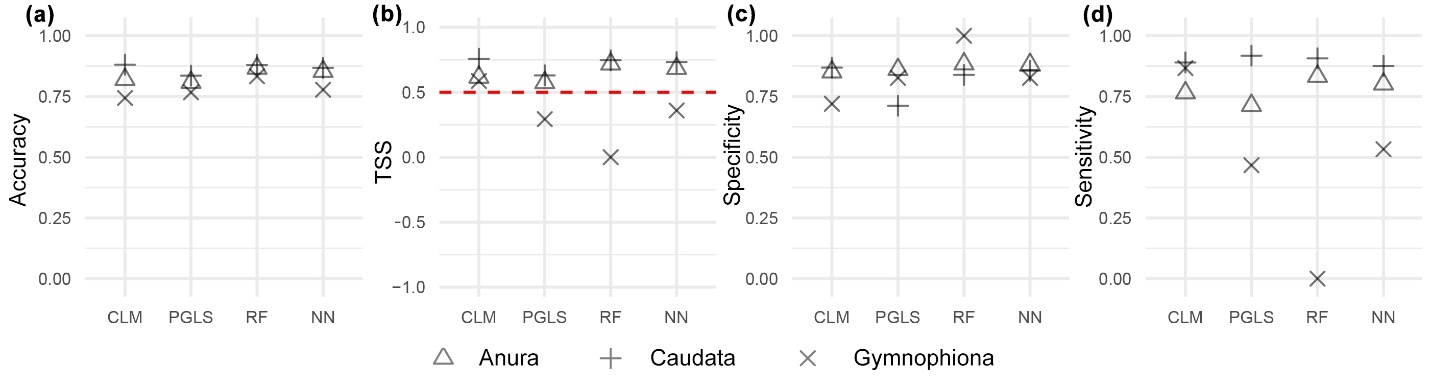

**Table A21.** Result for random block validation using as response variable Red List categories. We show the accuracy (rate of correct classification) the true skill statistic (TSS = sensitivity + specificity -1), the sensitivity, the specificity, and the mean absolute error (Error) for each category within each of the 12 fitted models (3 orders x 4 model algorithms).

| **Order** | **Algorithm** | **Category** | **Accuracy** | **TSS** | **Sensitivity** | **Specificity** | **Error** |
| --- | --- | --- | --- | --- | --- | --- | --- |
| ANURA | CLM | LC | 0.78 | 0.60 | 0.66 | 0.93 | 0.55 |
| ANURA | CLM | NT | 0.77 | 0.19 | 0.39 | 0.79 | 0.92 |
| ANURA | CLM | VU | 0.84 | 0.20 | 0.30 | 0.90 | 0.91 |
| ANURA | CLM | EN | 0.84 | 0.31 | 0.39 | 0.93 | 0.84 |
| ANURA | CLM | CR | 0.90 | 0.62 | 0.70 | 0.92 | 0.50 |
| ANURA | PGLS | LC | 0.68 | 0.43 | 0.46 | 0.97 | 0.67 |
| ANURA | PGLS | NT | 0.63 | 0.20 | 0.57 | 0.63 | 0.51 |
| ANURA | PGLS | VU | 0.73 | 0.16 | 0.39 | 0.77 | 0.65 |
| ANURA | PGLS | EN | 0.78 | 0.05 | 0.14 | 0.91 | 1.11 |
| ANURA | PGLS | CR | 0.92 | 0.03 | 0.04 | 1.00 | 1.51 |
| ANURA | RF | LC | 0.86 | 0.70 | 0.91 | 0.79 | 0.24 |
| ANURA | RF | NT | 0.94 | 0.14 | 0.14 | 1.00 | 1.16 |
| ANURA | RF | VU | 0.88 | 0.29 | 0.34 | 0.95 | 1.03 |
| ANURA | RF | EN | 0.86 | 0.63 | 0.73 | 0.89 | 0.49 |
| ANURA | RF | CR | 0.94 | 0.57 | 0.60 | 0.97 | 0.65 |
| ANURA | NN | LC | 0.81 | 0.65 | 0.73 | 0.93 | 0.38 |
| ANURA | NN | NT | 0.79 | 0.17 | 0.35 | 0.82 | 0.77 |
| ANURA | NN | VU | 0.81 | 0.28 | 0.43 | 0.85 | 0.68 |
| ANURA | NN | EN | 0.82 | 0.32 | 0.42 | 0.90 | 0.76 |
| ANURA | NN | CR | 0.92 | 0.21 | 0.23 | 0.99 | 1.06 |
| CAUDATA | CLM | LC | 0.86 | 0.62 | 0.67 | 0.95 | 0.44 |
| CAUDATA | CLM | NT | 0.83 | 0.34 | 0.48 | 0.86 | 0.61 |
| CAUDATA | CLM | VU | 0.78 | 0.27 | 0.41 | 0.86 | 0.69 |
| CAUDATA | CLM | EN | 0.79 | 0.37 | 0.48 | 0.89 | 0.60 |
| CAUDATA | CLM | CR | 0.94 | 0.86 | 0.91 | 0.95 | 0.09 |
| CAUDATA | PGLS | LC | 0.77 | 0.24 | 0.24 | 1.00 | 1.03 |
| CAUDATA | PGLS | NT | 0.74 | 0.25 | 0.48 | 0.76 | 0.68 |
| CAUDATA | PGLS | VU | 0.72 | 0.20 | 0.41 | 0.79 | 0.62 |
| CAUDATA | PGLS | EN | 0.75 | 0.33 | 0.49 | 0.84 | 0.55 |
| CAUDATA | PGLS | CR | 0.91 | 0.70 | 0.77 | 0.94 | 0.24 |
| CAUDATA | RF | LC | 0.86 | 0.70 | 0.83 | 0.88 | 0.34 |
| CAUDATA | RF | NT | 0.88 | 0.10 | 0.14 | 0.96 | 1.00 |
| CAUDATA | RF | VU | 0.83 | 0.41 | 0.51 | 0.90 | 0.68 |
| CAUDATA | RF | EN | 0.84 | 0.58 | 0.70 | 0.89 | 0.40 |
| CAUDATA | RF | CR | 0.95 | 0.82 | 0.86 | 0.97 | 0.17 |
| CAUDATA | NN | LC | 0.82 | 0.46 | 0.50 | 0.96 | 0.63 |
| CAUDATA | NN | NT | 0.76 | 0.29 | 0.50 | 0.79 | 0.59 |
| CAUDATA | NN | VU | 0.78 | 0.31 | 0.46 | 0.85 | 0.62 |
| CAUDATA | NN | EN | 0.75 | 0.30 | 0.45 | 0.85 | 0.66 |
| CAUDATA | NN | CR | 0.91 | 0.63 | 0.67 | 0.95 | 0.33 |
| GYMNOPHIONA | CLM | LC | 0.69 | 0.62 | 0.62 | 1.00 | 0.70 |
| GYMNOPHIONA | CLM | NT | 0.86 | -0.12 | 0.00 | 0.88 | 1.00 |
| GYMNOPHIONA | CLM | VU | 0.76 | 0.43 | 0.67 | 0.76 | 0.33 |
| GYMNOPHIONA | CLM | EN | 0.89 | 0.45 | 0.50 | 0.95 | 0.58 |
| GYMNOPHIONA | CLM | CR | 0.99 | NA | NA | 0.99 | NA |
| GYMNOPHIONA | PGLS | LC | 0.53 | 0.42 | 0.42 | 1.00 | 0.74 |
| GYMNOPHIONA | PGLS | NT | 0.57 | 0.07 | 0.50 | 0.57 | 0.50 |
| GYMNOPHIONA | PGLS | VU | 0.77 | -0.21 | 0.00 | 0.79 | 1.00 |
| GYMNOPHIONA | PGLS | EN | 0.89 | 0.17 | 0.17 | 1.00 | 1.25 |
| GYMNOPHIONA | PGLS | CR | 1.00 | NA | NA | 1.00 | NA |
| GYMNOPHIONA | RF | LC | 0.88 | 0.49 | 0.96 | 0.53 | 0.04 |
| GYMNOPHIONA | RF | NT | 0.84 | -0.14 | 0.00 | 0.86 | 1.00 |
| GYMNOPHIONA | RF | VU | 0.97 | 0.00 | 0.00 | 1.00 | 1.00 |
| GYMNOPHIONA | RF | EN | 0.87 | 0.00 | 0.00 | 1.00 | 2.50 |
| GYMNOPHIONA | RF | CR | 1.00 | NA | NA | 1.00 | NA |
| GYMNOPHIONA | NN | LC | 0.72 | 0.66 | 0.66 | 1.00 | 0.52 |
| GYMNOPHIONA | NN | NT | 0.77 | 0.27 | 0.50 | 0.77 | 0.50 |
| GYMNOPHIONA | NN | VU | 0.77 | 0.11 | 0.33 | 0.78 | 0.67 |
| GYMNOPHIONA | NN | EN | 0.86 | -0.01 | 0.00 | 0.99 | 1.42 |
| GYMNOPHIONA | NN | CR | 1.00 | NA | NA | 1.00 | NA |

**Table A22.** Results for averaged values among categories of random block validation using as response variable Red List categories (Table A21). We show the averaged values among categories for accuracy (rate of correct classification), the true skill statistic (TSS = sensitivity + specificity -1), the sensitivity, the specificity, and the mean absolute error (Error) for each of the 12 fitted models (3 orders x 4 model algorithms).

| **Order** | **Algorithm** | **Accuracy** | **TSS** | **Error** |
| --- | --- | --- | --- | --- |
| ANURA | CLM | 0.83 | 0.38 | 0.74 |
| ANURA | PGLS | 0.83 | 0.33 | 0.73 |
| ANURA | RF | 0.75 | 0.17 | 0.89 |
| ANURA | NN | 0.90 | 0.47 | 0.71 |
| CAUDATA | CLM | 0.84 | 0.49 | 0.49 |
| CAUDATA | PGLS | 0.80 | 0.40 | 0.57 |
| CAUDATA | RF | 0.78 | 0.34 | 0.62 |
| CAUDATA | NN | 0.87 | 0.52 | 0.52 |
| GYMNOPHIONA | CLM | 0.84 | 0.34 | 0.65 |
| GYMNOPHIONA | PGLS | 0.82 | 0.26 | 0.78 |
| GYMNOPHIONA | RF | 0.75 | 0.11 | 0.87 |
| GYMNOPHIONA | NN | 0.91 | 0.09 | 1.14 |

**Table A23.** Results averaged by the number of species for random block validation using as response variable Red List categories. We show the averaged values for accuracy (rate of correct classification), the true skill statistic (TSS = sensitivity + specificity -1), the sensitivity, the specificity, and the mean absolute error (Error) for each of the 12 fitted models (3 orders x 4 model algorithms).

| **Order** | **Algorithm** | **Accuracy** | **Error** | **TSS** |
| --- | --- | --- | --- | --- |
| ANURA | CLM | 0.56 | 0.66 | 0.48 |
| ANURA | PGLS | 0.37 | 0.80 | 0.29 |
| ANURA | RF | 0.74 | 0.46 | 0.60 |
| ANURA | NN | 0.57 | 0.56 | 0.49 |
| CAUDATA | CLM | 0.60 | 0.48 | 0.51 |
| CAUDATA | PGLS | 0.44 | 0.67 | 0.33 |
| CAUDATA | RF | 0.68 | 0.45 | 0.58 |
| CAUDATA | NN | 0.51 | 0.58 | 0.40 |
| GYMNOPHIONA | CLM | 0.59 | 0.68 | 0.57 |
| GYMNOPHIONA | PGLS | 0.38 | 0.81 | 0.36 |
| GYMNOPHIONA | RF | 0.78 | 0.42 | 0.39 |
| GYMNOPHIONA | NN | 0.56 | 0.64 | 0.54 |

**Table A24.** Results for random block validation using as response variable threatened/non threatened. We show the accuracy (rate of correct classification), the true skill statistic (TSS = sensitivity + specificity -1), the sensitivity and the specificity for each of the 12 fitted models (3 orders x 4 model algorithms).

| **Order** | **Algorithm** | **Accuracy** | **TSS** | **Sensitivity** | **Specificity** |
| --- | --- | --- | --- | --- | --- |
| ANURA | CLM | 0.82 | 0.62 | 0.76 | 0.85 |
| ANURA | PGLS | 0.81 | 0.57 | 0.71 | 0.86 |
| ANURA | RF | 0.87 | 0.72 | 0.83 | 0.88 |
| ANURA | NN | 0.85 | 0.68 | 0.80 | 0.88 |
| CAUDATA | CLM | 0.88 | 0.76 | 0.89 | 0.87 |
| CAUDATA | PGLS | 0.84 | 0.63 | 0.92 | 0.71 |
| CAUDATA | RF | 0.88 | 0.75 | 0.91 | 0.84 |
| CAUDATA | NN | 0.87 | 0.73 | 0.88 | 0.86 |
| GYMNOPHIONA | CLM | 0.74 | 0.59 | 0.87 | 0.72 |
| GYMNOPHIONA | PGLS | 0.77 | 0.29 | 0.47 | 0.83 |
| GYMNOPHIONA | RF | 0.83 | 0.00 | 0.00 | 1.00 |
| GYMNOPHIONA | NN | 0.78 | 0.36 | 0.53 | 0.83 |

**Figure A10.** Histograms showing (a) the difference between the ensemble prediction and the current IUCN RL category and (b) the *Species Prioritization Index* (*SPI*). Vertical lines with percentages in panel b describes the quantiles for the *SPI*.

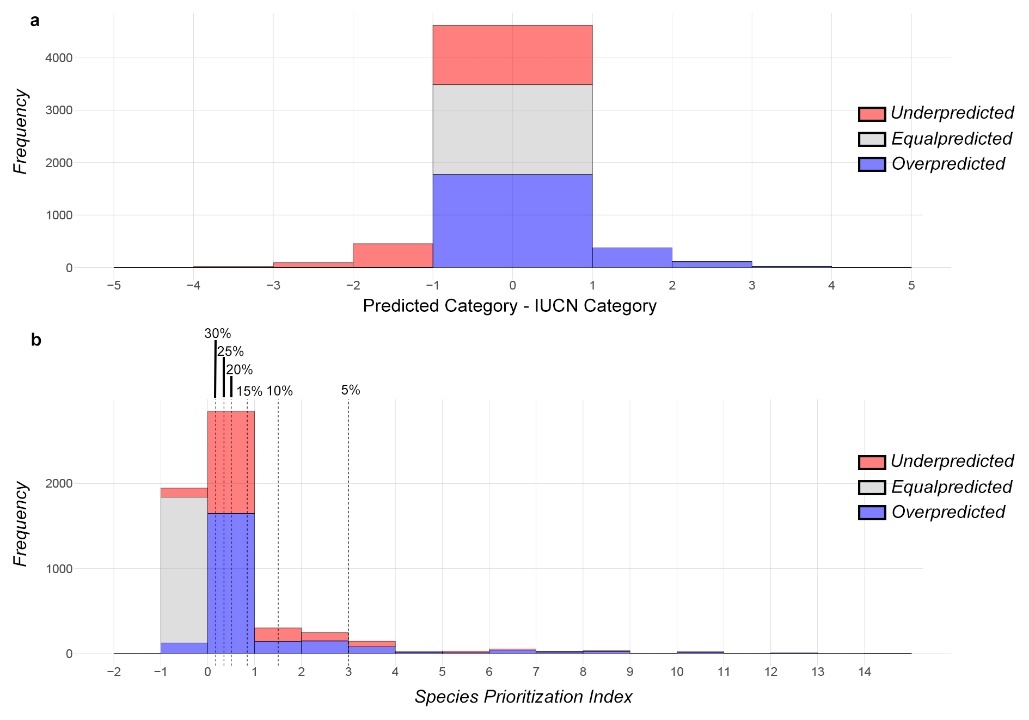

**Figure A11.** Maps showing the results for the (a) *Species Prioritization Index* *Underpredicted species* (*SPI_U_*), i.e., species whose Red List category is higher than predicted by the model. (b) *Species Prioritization Index* for *Overpredicted species* (*SPI_O_*), i.e. species with a IUCN Red List category lower than that predicted by the model. A, B and C on the map point to the location of the species A, B and C in Fig. 43, which are examples of *Underpredicted*, *Equally predicted* (species with a IUCN Red List category equal to that predicted by the model) and *Overpredicted* species.

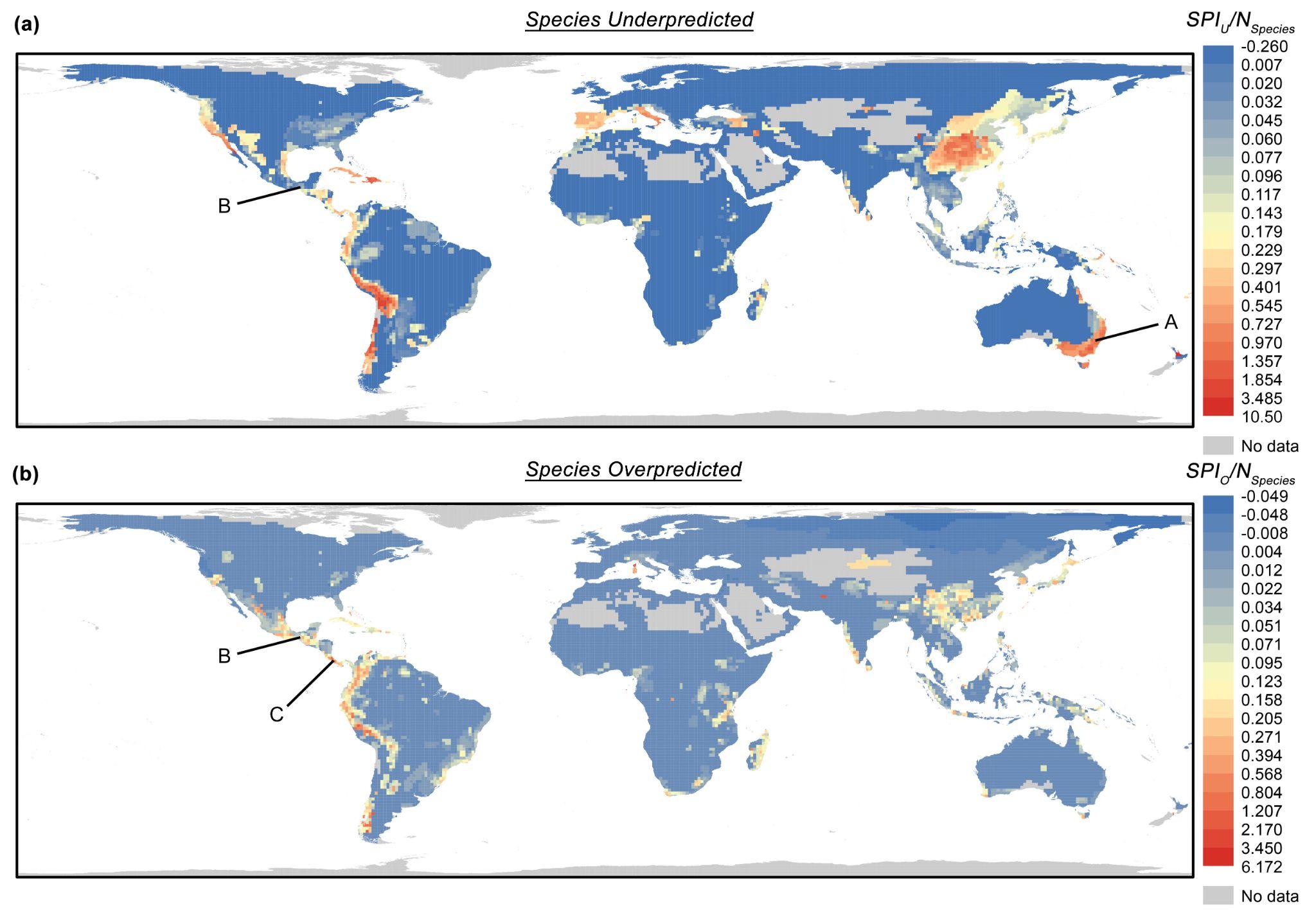

**Figure A12.** Range and median value by family for: (a) the *Species Prioritization Index* *Underpredicted* (*SPI_O_*) for *underpredicted species*, species which RL category is higher than our predicted category; and (b) the *Species Prioritization Index* *Overpredicted* (*SPI_O_*) for *overpredicted species*, species which RL category shows a lower threat category than our predicted category.

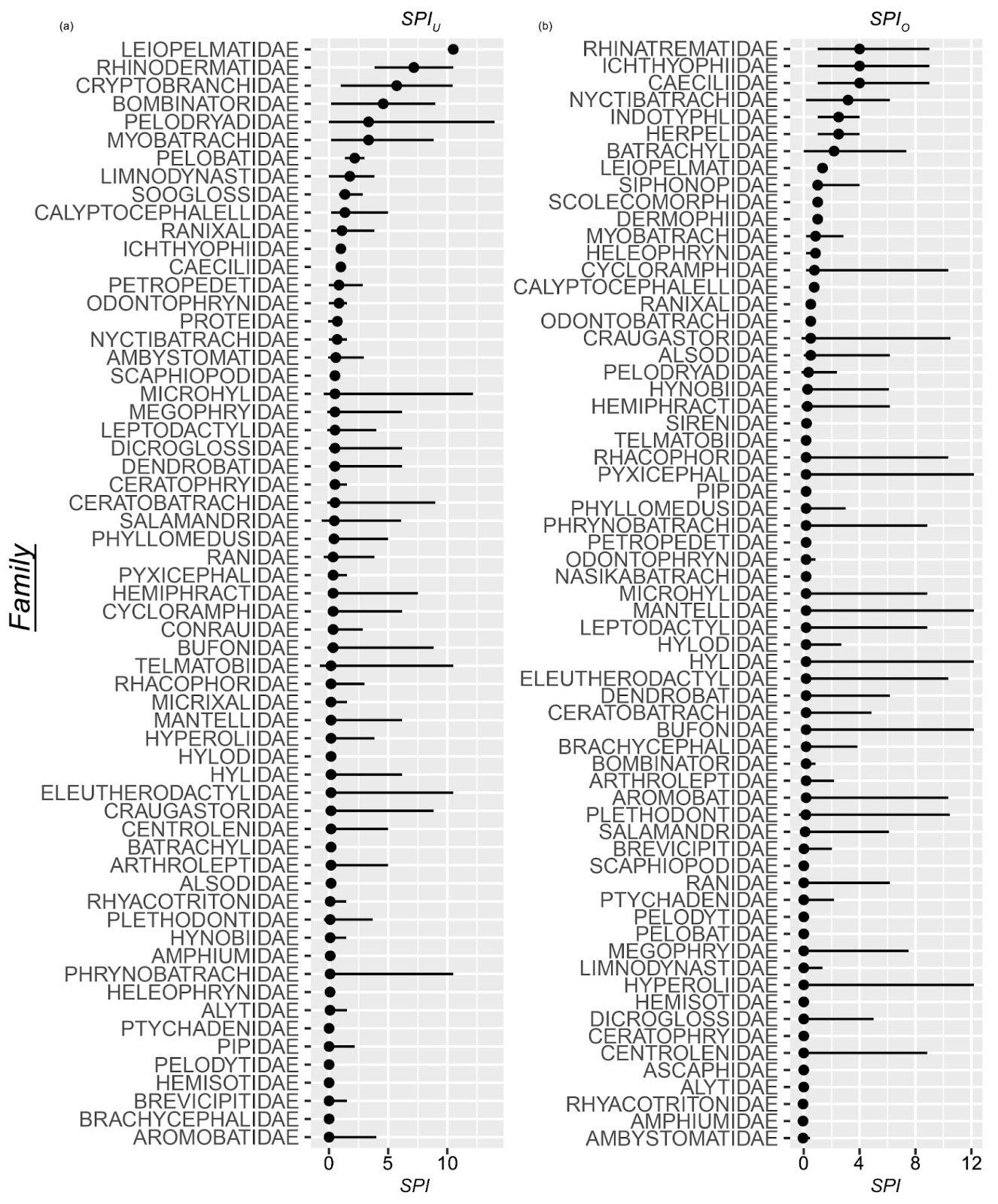

**REFERENCES**

Benítez-López, A., Santini, L., Schipper, A.M., Busana, M., Huijbregts, M.A.J., 2019. Intact but empty forests? Patterns of hunting-induced mammal defaunation in the tropics. PLoS Biology 17, e3000247.

Cardillo, M., 2021. Clarifying the relationship between body size and extinction risk in amphibians by complete mapping of model space. Proceedings of the Royal Society B: Biological Sciences 288, 20203011.

Cardillo, M., Mace, G.M., Gittleman, J.L., Jones, K.E., Bielby, J., Purvis, A., 2008. The predictability of extinction: biological and external correlates of decline in mammals. Proceedings of the Royal Society of London, Series B: Biological Sciences 275, 1441-1448.

Carilo Filho, L.M., de Carvalho, B.T., Azevedo, B.K.A., Gutiérrez-Pesquera, L.M., Mira-Mendes, C.V., Solé, M., Orrico, V.G.D., 2021. Natural history predicts patterns of thermal vulnerability in amphibians from the Atlantic Rainforest of Brazil. Ecology and evolution 11, 16462-16472.

Christensen, R.H.B., 2020. Ordinal package version 2020.8-22.

ESA, 2021. Land cover classification gridded maps from 1992 to present derived from satellite observations version 2.0 and 2.1.

Evans, J., Murphy, M., 2018. rfUtilities. R package version 2.1-3.

Ficetola, G.F., Rondinini, C., Bonardi, A., Baisero, D., Padoa-Schioppa, E., 2015. Habitat availability for amphibians and extinction threat: a global analysis. Diversity and Distributions 21, 302-311.

González-del-Pliego, P., Freckleton, R.P., Edwards, D.P., Koo, M.S., Scheffers, B.R., Pyron, R.A., Jetz, W., 2019. Phylogenetic and Trait-Based Prediction of Extinction Risk for Data-Deficient Amphibians. Current Biology 29, 1557-1563.e1553.

Hijmans, R.J., Phillips, S., Leathwick , J., Elith, J., 2020. dismo: Species Distribution Modeling. R package version 1.3-3.

IUCN, 2021. IUCN Red List of Threatened Species Version 2021-2

Jetz, W., Pyron, R.A., 2018. The interplay of past diversification and evolutionary isolation with present imperilment across the amphibian tree of life. Nature Ecology & Evolution 2, 850-858.

Karger, D.N., Conrad, O., Böhner, J., Kawohl, T., Kreft, H., Soria-Auza, R.W., Zimmermann, N.E., Linder, H.P., Kessler, M., 2017. Climatologies at high resolution for the earth’s land surface areas. Scientific data 4, 170122.

Karger, D.N., Conrad, O., Böhner, J., Kawohl, T., Kreft, H., Soria-Auza, R.W., Zimmermann, N.E., Linder, H.P., Kessler, M., 2018. Data from: Climatologies at high resolution for the earth's land surface areas, Dryad, Dataset.

LeDell, E., Gill, N., Aiello, S., Fu, A., Candel, A., Click, C., Kraljevic, T., Nykodym, T., Aboyoun, P., Kurka, M., Malohlava, M., 2022. h2o: R Interface for the 'H2O' Scalable Machine Learning Platform. R package version 3.38.0.1.

Liaw, A., Wiener, M., 2002. Classification and Regression by randomForest. R News 2, 18-22.

Lucas, P.M., González‐Suárez, M., Revilla, E., 2019. Range area matters, and so does spatial configuration: predicting conservation status in vertebrates. Ecography.

McKinney, M.L., 2002. Urbanization, Biodiversity, and Conservation. Bioscience 52, No. 10.

Murphy, M.A., Evans, J.S., Storfer, A., 2010. Quantifying Bufo boreas connectivity in Yellowstone National Park with landscape genetics. Ecology 91, 252-261.

NASA, 2018. Gridded Population of the World, Version 4 (GPWv4): Population Density, Revision 11. NASA Socioeconomic Data and Applications Center (SEDAC), Palisades, NY.

Newbold, T., Hudson, L.N., Hill, S.L.L., Contu, S., Lysenko, I., Senior, R.A., Borger, L., Bennett, D.J., Choimes, A., Collen, B., Day, J., De Palma, A., Diaz, S., Echeverria-Londono, S., Edgar, M.J., Feldman, A., Garon, M., Harrison, M.L.K., Alhusseini, T., Ingram, D.J., Itescu, Y., Kattge, J., Kemp, V., Kirkpatrick, L., Kleyer, M., Correia, D.L.P., Martin, C.D., Meiri, S., Novosolov, M., Pan, Y., Phillips, H.R.P., Purves, D.W., Robinson, A., Simpson, J., Tuck, S.L., Weiher, E., White, H.J., Ewers, R.M., Mace, G.M., Scharlemann, J.P.W., Purvis, A., 2015. Global effects of land use on local terrestrial biodiversity. Nature 520, 45-50.

Nowakowski, A.J., Thompson, M.E., Donnelly, M.A., Todd, B.D., 2017. Amphibian sensitivity to habitat modification is associated with population trends and species traits. Global Ecology and Biogeography 26, 700-712.

Oliveira, B.F., São-Pedro, V.A., Santos-Barrera, G., Penone, C., Costa, G.C., 2017. AmphiBIO, a global database for amphibian ecological traits. Scientific data 4, 170123.

Olson, D.M., Dinerstein, E., Wikramanayake, E.D., Burgess, N.D., Powell, G.V.N., Underwood, E.C., D'amico, J.A., Itoua, I., Strand, H.E., Morrison, J.C., Loucks, C.J., Allnutt, T.F., Ricketts, T.H., Kura, Y., Lamoreux, J.F., Wettengel, W.W., Hedao, P., Kassem, K.R., 2001. Terrestrial Ecoregions of the World: A New Map of Life on Earth. Bioscience 51, 933-938, 936.

Paradis, E., Schliep, K., 2019. ape 5.0: an environment for modern phylogenetics and evolutionary analyses in R. Bioinformatics (Oxford) 35, 526-528.

Pincheira-Donoso, D., Harvey, L.P., Cotter, S.C., Stark, G., Meiri, S., Hodgson, D.J., 2021a. The global macroecology of brood size in amphibians reveals a predisposition of low-fecundity species to extinction. Global Ecology and Biogeography 30, 1299-1310.

Pincheira-Donoso, D., Harvey, L.P., Grattarola, F., Jara, M., Cotter, S.C., Tregenza, T., Hodgson, D.J., 2021b. The multiple origins of sexual size dimorphism in global amphibians. Global Ecology and Biogeography 30, 443-458.

Pinheiro, J., Bates, D., DebRoy, S., Sarkar, D., Team, R.C., 2021. Linear and Nonlinear Mixed Effects Models. R package version 3.1-153.

Skelly, D.K., Werner, E.E., Cortwright, S.A., 1999. Long-term distributional dynamics of a Michigan amphibian assemblage. Ecology 80, 2326-2337.

Venables, W.N., Ripley, B.D., 2002. Modern Applied Statistics with S, Fourth edn. Springer, New York.

Weiss, D.J., Nelson, A., Gibson, H.S., Temperley, W., Peedell, S., Lieber, A., Hancher, M., Poyart, E., Belchior, S., Fullman, N., Mappin, B., Dalrymple, U., Rozier, J., Lucas, T.C.D., Howes, R.E., Tusting, L.S., Kang, S.Y., Cameron, E., Bisanzio, D., Battle, K.E., Bhatt, S., Gething, P.W., 2018. A global map of travel time to cities to assess inequalities in accessibility in 2015. Nature 553, 333-336.

Yackulic, C.B., Sanderson, E.W., Uriarte, M., 2011. Anthropogenic and environmental drivers of modern range loss in large mammals. Proceedings of the National Academy of Sciences USA 108, 4024-4029.
